# Intragenic repeat expansions control yeast chronological aging

**DOI:** 10.1101/653006

**Authors:** Benjamin P Barré, Johan Hallin, Jia-Xing Yue, Karl Persson, Ekaterina Mikhalev, Agurtzane Irizar, Dawn Thompson, Mikael Molin, Jonas Warringer, Gianni Liti

## Abstract

Aging varies among individuals due to both genetics and environment but the underlying molecular mechanisms remain largely unknown. Using a highly recombined *Saccharomyces cerevisiae* population, we found 30 distinct Quantitative Trait Loci (QTLs) that control chronological life span (CLS) in calorie rich and calorie restricted environments, and under rapamycin exposure. Calorie restriction and rapamycin extended life span in virtually all genotypes, but through different genetic variants. We tracked the two major QTLs to massive expansions of intragenic tandem repeats in the cell wall glycoproteins *FLO11* and *HPF1*, which caused a dramatic life span shortening. Life span impairment by N-terminal *HPF1* repeat expansion was partially buffered by rapamycin but not by calorie restriction. The *HPF1* repeat expansion shifted yeast cells from a sedentary to a buoyant state, thereby increasing their exposure to surrounding oxygen. The higher oxygenation perturbed methionine, lipid, and purine metabolism, which likely explains the life span shortening. We conclude that fast evolving intragenic repeat expansions can fundamentally change the relationship between cells and their environment with profound effects on cellular life style and longevity.

## INTRODUCTION

Aging is a progressive decline in biological functions occurring in almost all living organisms that ultimately leads to death (Finch 1990; Jones et al. 2014). The first life span regulating genes were identified in the beginning of the 90s (Johnson 1990; Kenyon et al. 1993; Sun et al. 1994). Today hundreds have been uncovered (Kenyon 2010), although most are of small effect and few explain aging variation between individuals. Besides genetics, environmental factors, such as calorie restriction (CR) (Colman et al. 2009; Jiang 2000; Klass 1977; Pletcher et al. 2002; Weindruch et al. 1986), reduced oxygen exposure (Leiser et al. 2013; Rascon and Harrison 2010), and low temperature (Conti et al. 2006; Leiser, Begun, and Kaeberlein 2011; Sestini, Carlson, and Allsopp 1991), extend longevity. How genetics and environment interact to control variation in life span and by which mechanisms remains poorly understood. The beneficial effect of calorie restriction on longevity in organisms ranging from yeast (Lin, Defossez, and Guarente 2000) to primates (Mattison et al. 2017) has been known for >80 years (McCay, Crowell, and Maynard 1935) and is still the most successful intervention to delay aging, although its impact on life span has sometimes been disputed (Liao et al. 2010; Schleit et al. 2013). Cellular mediation of CR is at least in part occurring through nutrient sensitive signalling networks, including the insulin/IGF-1, mTOR (target of rapamycin), cAMP-PKA and AMPK pathways. These are held to regulate life span by controlling stress responses, mitochondrial respiration, redox homeostasis, genome stability, autophagy, energy and fat metabolism (Alvers et al. 2009; Hansen et al. 2008; Madia et al. 2008; Molin et al. 2011; Schulz et al. 2007; Wei et al. 2008; Weinberger et al. 2007; Yuan et al. 2012). Pharmaceutical control of some of these pathways can extend longevity in model organisms. Rapamycin, a clinically approved TOR inhibitor (Eisenberg et al. 2009; De Haes et al. 2014; Harrison et al. 2009; Martin-Montalvo et al. 2013), extends life span by mimicking CR (Blagosklonny 2010), but undesirable side effects in humans restrict its usage (Kaeberlein 2014).

The budding yeast *S. cerevisiae* has been pivotal in elucidating mechanisms regulating aging. Yeast aging can be studied through two approaches: replicative life span (RLS) and chronological life span (CLS). Replicative life span is the number of mitotic divisions before senescence and is used as a paradigm to study aging of proliferative tissues, such as stem cells (Mortimer and Johnston 1959; Steinkraus, Kaeberlein, and Kennedy 2008). Chronological lifespan is the time yeasts survive in non-proliferative conditions and models the aging of post-mitotic cells, such as neurons (Longo et al. 2012; Longo, Gralla, and Valentine 1996). Hundreds of genes whose disruption affect the CLS of lab domesticated yeast in calorie rich (Fabrizio et al. 2010; Garay et al. 2014; Powers et al. 2006) and calorie-restricted (Matecic et al. 2010) environments have been identified. However, most studies relied on artificial gene deletions and were performed in lab domesticated strains, which are highly atypical (Warringer et al. 2011), maintained as haploids rather than diploids (Peter et al. 2018), carry auxotrophies that alter life span (Boer, Amini, and Botstein 2008; Gomes et al. 2007), and have never been exposed to natural selection. Thus, genetic variants that regulate natural life span variation are still largely unknown. Crosses between natural yeast strains have the potential to uncover these variants (Brem 2002; Steinmetz et al. 2002), but remain poorly explored. Previous work linked natural polymorphisms in the ribosomal DNA and in the sirtuin *SIR2* (Kwan et al. 2013; Stumpferl et al. 2012), as well as telomere maintenance (Kwan et al. 2011) and serine biosynthesis (Jung et al. 2018) to life span variation. Lack of genetic diversity, mapping resolution and power has prevented more exhaustive exploration.

We unravelled the genetic basis of CLS variation using advanced intercrosses between two natural *S. cerevisiae* isolates with very different life spans. We measured CLS in calorie rich and calorie restricted media, and under rapamycin treatment. We mapped 30 unique QTLs of which the two major were explained by massive intragenic tandem repeat expansions in the cell wall glycoproteins *FLO11* and *HPF1* that dramatically shortened life span. Serine/threonine repeat expansions close to the *HPF1* N-terminus shifted cells from a sedentary to a buoyant life style, thereby increasing exposure to oxygen and perturbing methionine, lipid and purine homeostasis. Interestingly, the downstream effects of buoyancy were partially recovered by rapamycin treatment, suggesting that they can be targeted by TORC1 repression.

## RESULTS

### Calorie restriction and rapamycin extend life span through different genetic variants

We crossed a long-lived North American (NA) oak tree bark strain (YPS128) with a short-lived West African (WA) palm wine strain (DBVPG6044) which differ at 0.53% of nucleotide sites (Liti et al. 2009; Parts et al. 2011). A pool of F12 segregants of opposite mating types were then mated to generate 1056 diploids with hybrid, phased genomes, termed Phased Outbred Lines (POLs) (Hallin et al. 2016). POLs were individually cultivated in calorie rich (synthetic dextrose complete, SDC), calorie restricted (CR), or rapamycin supplemented (RM) environments for the whole experiment, and viability was measured by high throughput flow cytometry at 7, 21, and 35 days after media exhaustion. A total of 52466 genetic markers were called and used to run a genome-wide linkage analysis.

We found a remarkable lifespan diversity (Fig. 1A and table S3), with survival rates ranging from 6 to 97% already after 7 days in SDC (47% mean viability). Calorie restriction (86% mean viability at day 7) and RM (83% mean viability at day 7) sharply extended life spans of all genotypes (with a single exception in RM) (Fig. 1A and S1A). Life span was fairly correlated across environments (Pearson’s *r* = 0.62 for SDC vs CR, 0.53 for SDC vs RM; Fig. 1B) implying that CLS is mainly regulated by shared genetic effects across environments. Nevertheless, the more modest correlation between CR and RM (Pearson’s *r* = 0.43; Fig. 1C) suggested that partially distinct molecular mechanisms controlled their life span extension effect.

**Fig 1.**
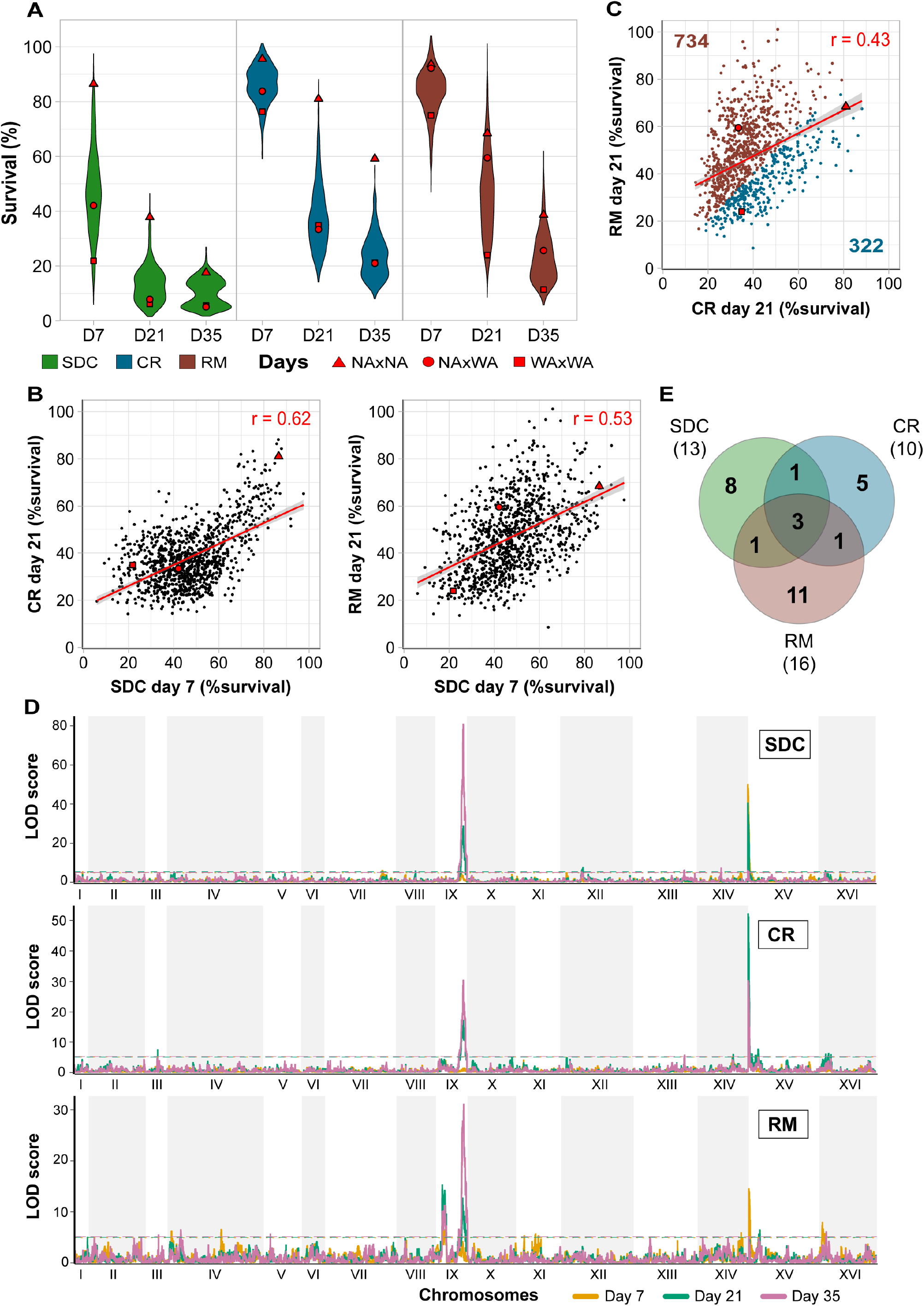
Calorie restriction and rapamycin extend life span through different genetic variants. Chronological life span of 1056 diploid segregant lineages from an F12 NA/WA advanced intercross. CLS was measured by counting viable cells (%) 7, 21 and 35 days after entry into quiescence, following growth in calorie sufficient (SDC), restricted (CR) and rapamycin (RM) media. Red: Founder homozygote parents (NA/NA, WA/WA) and their F1 hybrid (NA/WA). **(A)** CLS distributions. **(B)** Comparing CLS across environments and time points. Red line: linear regression, with 95% confidence interval. **(C)** CLS distributions. Numbers: lineages living longer in one environment. Blue: living longer in CR, brown: living longer in RM. **(D)** Linkage analysis of CLS. Panels: calorie rich (top), restricted (middle) and rapamycin (bottom) media. Line colour: 7 (yellow), 21 (green) and 35 (purple) days after entry into quiescence. *y*-axis: LOD score, *x*-axis: genome position. Dashed lines: significance QTL (α = 0.05). **(E)** QTLs private to and shared between environments. Numbers in parentheses: total QTLs per environment.

We found a total of 30 unique QTLs associated with chronological aging (Fig. 1D, S1B) that explained up to 40% of life span variation (Table S4). QTLs were mostly private to one environment and only three were detected in all (Fig. 1E). Two of these, located on chrIX and chrXV, were strikingly stronger than others and explained up to ~30% and ~20% of life span variation respectively (Table S4). Although both major QTLs were ubiquitous, the chrXV QTL was partially masked by RM treatment, whereas the chrIX QTL only became significant at advanced age (days 21 and 35) (table S4 and Fig. 1D and S1B). Most of the remaining QTLs were time and environment dependent and explained much less (mean: ~3%) of the life span variation (Table S4). Thus, CLS was largely determined by a few, very strong QTLs that were shared across calorie rich and calorie restricted environments. Chronological life span was then fine-tuned by mechanisms private to each environment; although the conservative threshold for calling QTLs may lead us to somewhat underestimate the shared QTLs.

### Natural variations in cell wall glycoproteins *HPF1* and *FLO11* control chronological life span

The two major QTLs peaked within *FLO11* (chrIX) and *HPF1* (chrXV). Both encode secreted cell wall glycoproteins with no known connections to life span. *Hpf1p* is functionally uncharacterized, while *Flo11p* regulates cell adhesion, pseudohyphae and biofilm formation (Douglas et al. 2007; Guo et al. 2000; Váchová et al. 2011). We found shorter life span for WA-*FLO11* and *HPF1* compared to NA homozygotes, with the WA short life span alleles being completely dominant (Fig. 2A). We validated these effects in a reciprocal hemizygosity assay (Steinmetz et al. 2002) (Fig. 2B and methods); a NA/WA hybrid deleted for the WA-*HPF1* allele lived 70% longer, regardless of the presence of NA-*HPF1* (Fig. 2C). Life span extension by rapamycin, but not by calorie restriction, rescued the WA-*HPF1* shortening, consistent with the QTL being exclusively weaker in rapamycin (Fig. 2C and 1D). The WA-*FLO11* also shortened life span but to a lesser extent compared to expectations from the QTL strength, perhaps due to either linkage or epistasis with other variants. As for *HPF1*, removing the WA-*FLO11* or both the WA and NA-*FLO11* alleles extended life span, while removing the NA-*FLO11* had no effect (Fig. 2D). The negative effect of WA-*FLO11* increased with age and was not rescued by rapamycin, again as expected (Fig. 2D and 1D). Removing both alleles of either *HPF1* or *FLO11* extended the life span of WA/WA but not NA/NA homozygote, and failed to affect the domesticated reference strain, S288C (Fig. S2).

**Fig 2.**
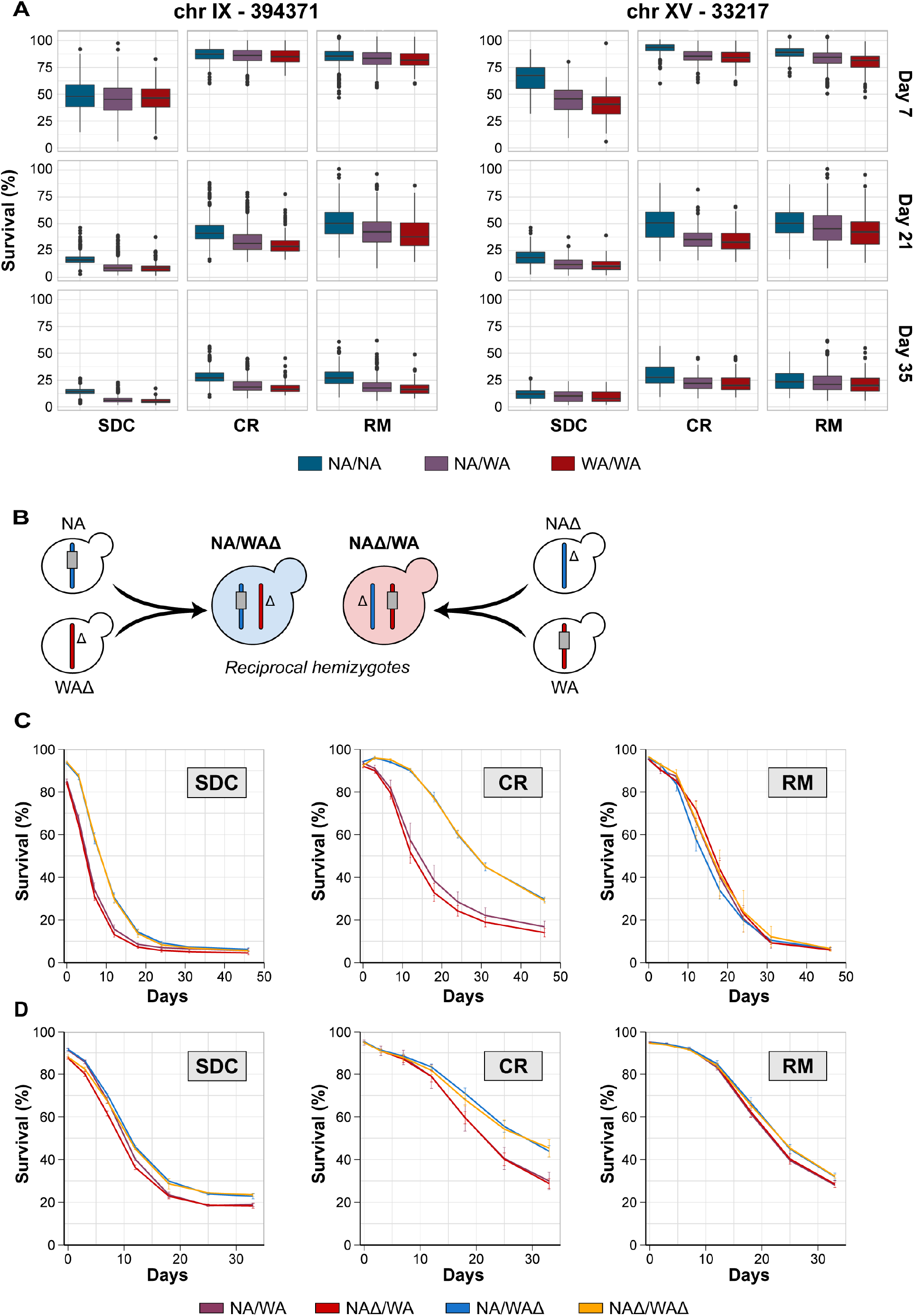
Natural variations in cell wall glycoproteins *HPF1* and *FLO11* control chronological life span. **(A)** Chronological life span of the 1056 segregant lineages from the F12 NA/WA advanced intercross. Lineages were separated according to genotype at the markers with highest LOD score in each of the two major QTLs: 394,381 kb in chromosome IX (in *FLO11*) and 33,217 kb in chromosome XV (in *HPF1*). **(B)** Schematic representation of the NA/WA reciprocal hemizygosity design used to validate the CLS effect of the *HPF1* and *FLO11* WA alleles. Colour: NA (blue) and WA (red) chromosomes. Grey rectangle: candidate gene (*HPF1, FLO11*). Δ: gene deletion. **(C)** Reciprocal hemizygosity. CLS of NA/WA hemizygotes for NA (blue; WAΔ) and WA (red; NAΔ) *HPF1*, heterozygote for *HPF1* (purple; NA/WA) and lacking *HPF1* (yellow; NAΔWAΔ). **(D)** As in C) but for *FLO11*.

### Massive intragenic tandem repeat expansions within *FLO11* and *HPF1* shorten life span

Remarkably, *FLO11* and *HPF1* both carry intragenic tandem repeats that are massively expanded in the WA allele. WA-*HPF1* is twice as long (6006 vs. 3033bp) and WA-*FLO11* is 10% longer (4014 vs. 3654bp) than their NA counterparts (Fig. 3A and S3A). Repeat motifs were between 21 and 71 amino acids long and mainly composed of threonine and serine, as reported for alleles of other cell wall proteins (Verstrepen and Klis 2006) (Fig. 3A). The partial motif degeneration in these repeat motifs prevented pinpointing their exact patterns and boundaries for both *HPF1* and *FLO11*. Such repeat expansion in WA alleles is gene specific; of 26 genes containing very long tandem repeats (Verstrepen et al. 2005) only *HPF1* was massively expanded in WA relative to six other strains for which complete genome assemblies exists (Yue et al. 2017) (Fig. 3B).

**Fig 3.**
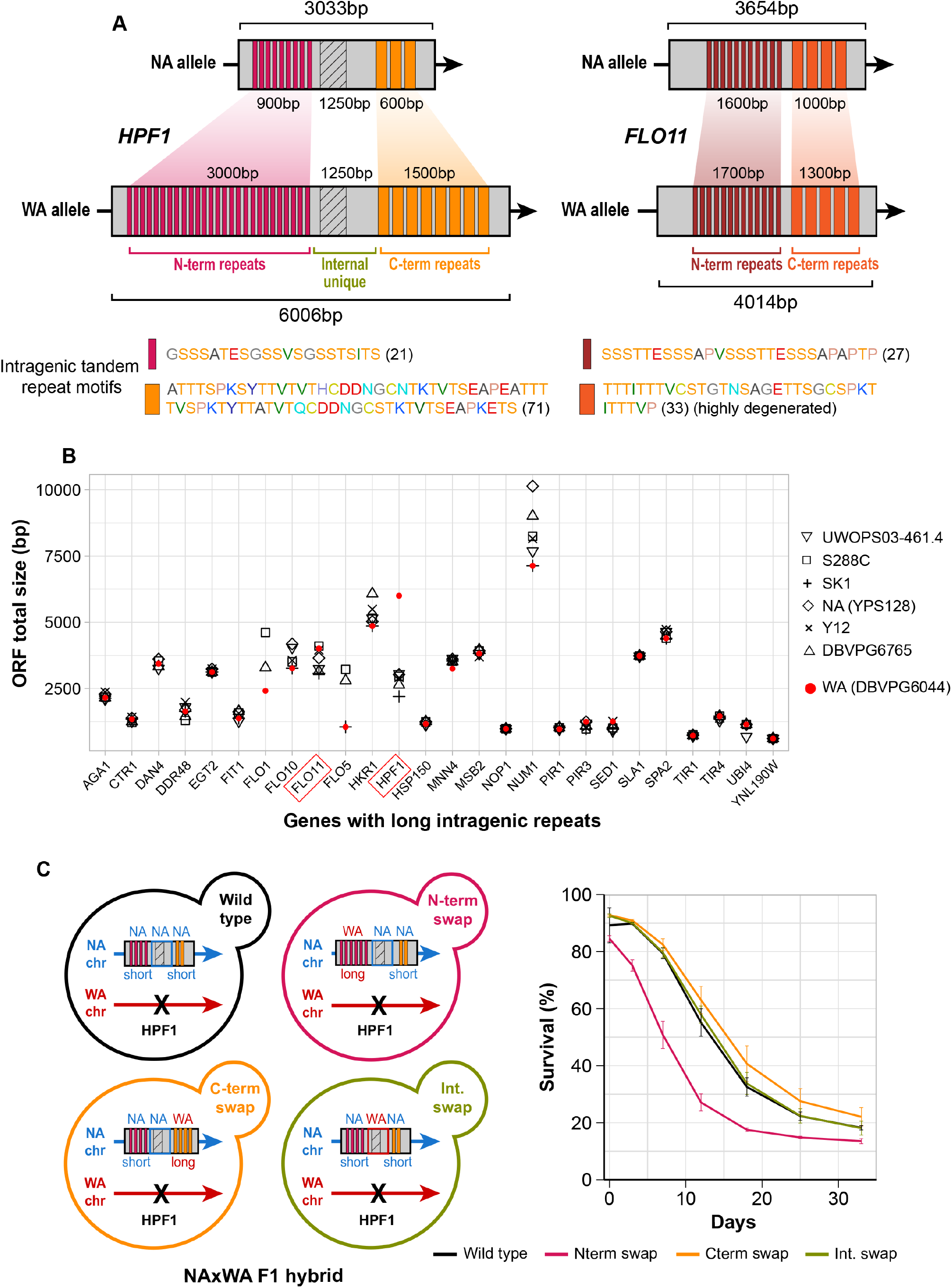
Massive intragenic tandem repeat expansions within *FLO11* and *HPF1* shorten life span. **(A)** Schematic representation of the intragenic repeats (coloured rectangles) in *HPF1* and *FLO11* in NA and WA. Hatched rectangle: a *HPF1* internal unique region with high sequence variation between NA and WA alleles. Zoom-in: repeat motifs. Amino acids are colored according to the RasMol nomenclature. Numbers = motif size (amino acids). **(B)** Size variation of genes containing long intragenic repeats in seven diverged *S. cerevisiae* strains (Verstrepen et al. 2005; Yue et al. 2017). Diamonds: North American. Red circles: West African. **(C)** Left panel: Design of allele swaps of *HPF1* segments in the NA/WA F1 hybrid. The WA-*HPF1* allele was deleted (black cross), while the NA-*HPF1* was kept unchanged (wildtype), or a segment was replaced by the corresponding WA-*HPF1* segment. N-term: N-terminal repeats, C-term: C-terminal repeats, Int: internal unique region. Right panel: CLS for allele swapped constructs.

We hypothesized that the WA-*HPF1* massive repeat expansions explained the life span shortening and tested this by swapping *HPF1* alleles in the F1 NA/WA hybrid. We removed WA-*HPF1*, while the remaining NA allele was engineered to contain specific segments of the WA allele (see methods). Swapped segments corresponded to N- and C-terminal blocks of tandem repeats, and to the highly polymorphic internal unique domain (Fig. 3A and 3C). Substituting the C-terminal repeats or the internal domain of NA-*HPF1* with its WA counterpart did not shorten CLS, but replacing the N-terminal repeats shortened CLS as much as the native WA allele (Fig. 3C). Likewise, inserting the WA N-terminal repeats into a NA homozygote diploid dramatically shortened life span (Fig. S3B), while inserting the NA N-terminal repeats into a WA homozygote diploid extended its life span to a comparable extent (Fig. S3C).

### Buoyancy triggered by *HPF1* N-terminal repeat expansions shortens life span

Intriguingly, NA/WA hybrids carrying only the WA-*HPF1* became buoyant during exponential growth, i.e. they shifted to a free-floating life style. Following entry into stationary phase, cells sedimented again, returning to a sedentary life style (Fig. 4A). We tracked this life style shift to the WA-*HPF1* N-terminal repeat expansion, by showing that it induces flotation (Fig. 4B). To probe whether buoyancy *per se* shortened lifespan, we repeated the CLS assay in intensely shaken flasks rather than in static 96 well-plates. Because intense shaking forces all yeast cells to remain in suspension, we postulated that it would eliminate the *HPF1* life span shortening only if it was due to buoyancy. In line with this assumption, we found that buoyancy enforced by shaking reduced life span dramatically and completely negated the effect of *HPF1* allelic variation on life span (Fig. 4C). Shorter life span in shaking cultures has previously been explained as a result of higher exposure to oxygen (Fabrizio et al. 2003; Longo et al. 1999). Thus, by promoting cellular buoyancy in static cultures, *HPF1* exposes cells to higher oxygen levels and shortens life span, without affecting medium acidity and with no effect of Hpf1p secretion to the medium (Fig. S4A and S4B). Complete *HPF1* removal in the WA homozygote nullified cell buoyancy and increased CLS, whereas it neither affected buoyancy nor life span in the NA homozygote (Fig. 4D). We conclude that buoyancy shortens life span and that *HPF1* controls life span by shifting cells between buoyant and sedentary life styles.

**Fig 4.**
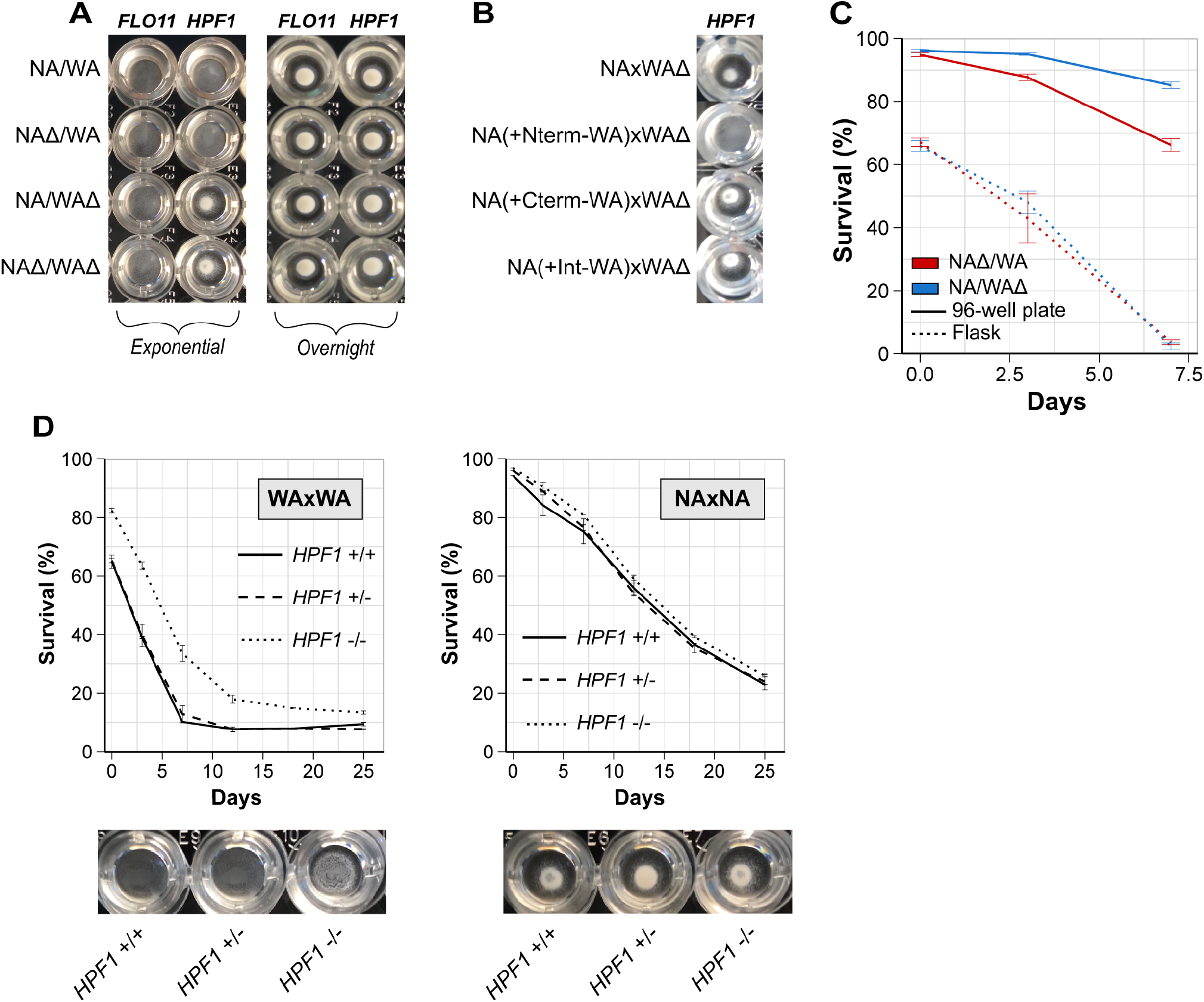
Buoyancy triggered by *HPF1* N-terminal repeat expansions shortens life span. **(A-B)** Buoyancy of cells cultivated for 7 hours (exponential phase) or overnight in calorie rich (SDC) medium in a 96-well plate. **A)** *HPF1* and *FLO1* hemizygotes. **B)** *HPF1* allele swaps (see Fig 3C). **(C)** Comparing CLS of *HPF1* hemizygote cells cultivated in shake flasks and 96-well plates. Shake flasks had a 5:1 volume/medium ratio and were shaken at 220rpm. 96-well plates were filled with 200 µL medium, with no shaking. **(D)** CLS (top panel) and buoyancy (96 well plates; exponential phase) of WA and NA homozygotes with no (full line), 1 (dashed lines) or both copies (dotted lines) of *HPF1* deleted in calorie rich medium (SDC).

### *HPF1* induced buoyancy reprograms methionine, lipid, and purine metabolism

We hypothesized that meeting the oxygen challenge associated with *HPF1* induced buoyancy would reprogram metabolism and gene expression. We therefore compared the transcriptomes of the two *HPF1* reciprocal hemizygous hybrids, before (exponential growth) and after (7 days) the onset of aging (Table S5), with or without rapamycin exposure. During exponential growth, buoyancy and higher oxygen exposure induced by the WA-*HPF1* only weakly affected transcript abundances (8 and 3 genes changing >2-fold in calorie rich and rapamycin media respectively, Fig. 5A and 5B). The known low oxygen responders *TIR1* and *ANB1* (Cohen et al. 2001; Lowry and Lieber 1986) were repressed by oxygen as expected. Five of the six transcripts induced by oxygenation encode methionine metabolic proteins (Fig. 5A), notably Mxr1p, reducing oxidised methionine. Induction of methionine reduction may signal increased methionine oxidation from ROS and explain the life span shortening. Rapamycin supplementation completely nullified the induction of methionine metabolism and reduction, despite cells being buoyant, potentially explaining why rapamycin prevents the WA-*HPF1* from shortening lifespan (Fig. 5A and S5A).

**Fig 5.**
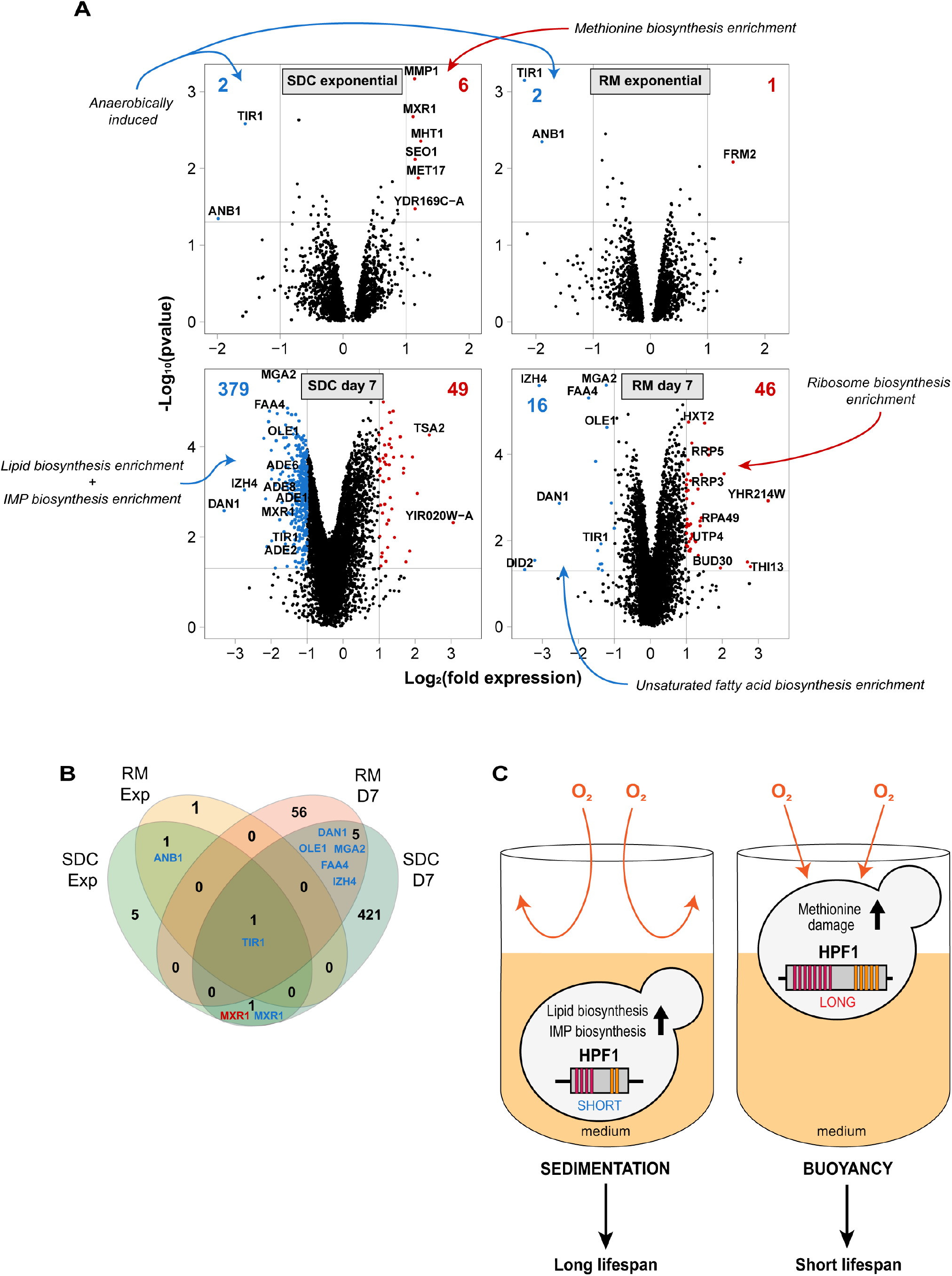
*HPF1* induced buoyancy reprograms methionine, lipid, and purine metabolism. **(A)** Transcriptome changes induced by WA-*HPF1* dependent buoyancy. NA/WA hybrids hemizygotes for WA or NA-*HPF1* were cultivated in calorie rich (left panels; SDC) or rapamycin (right panels; RM) medium and RNA was extracted and sequenced from exponential phase (top panels) or aging (bottom panels; 7 days after entry into quiescence) cells. *y*-axis:-log_10_ (*p*-value). *x*-axis: log_2_ (*HPF1* NAΔ/WA) -log_2_ (*HPF1* NA/WAΔ). Blue: transcripts more (>2x, p<0.05) abundant in *HPF1* NA/WAΔ. Red: transcripts more (>2x, p<0.05) abundant in *HPF1* NAΔ/WA. The total number of transcripts passing each criterion are reported (top corners, blue and red text). Gene ontology classifications enriched among transcripts passing each criterion are indicated (blue and red arrows, table S5). **(B)** Comparing number of transcripts more (>2x, p<0.05) abundant in *HPF1* NA/WAΔ (blue) or in *HPF1* NAΔ/WA (red) across environments (SDC and RM, in exponential phase and after 7 days of aging (D7)). **(C)** Model for how WA-*HPF1* shortens life span by shifting cells from a sedentary to a buoyant life style, exposing them to more oxygen, repressing lipid and purine biosynthesis and oxidizing free and protein bound methionine.

In sharp contrast to before the onset of aging, the oxygen exposure induced by buoyancy fundamentally (428 and 62 genes changing >2-fold, in calorie rich and rapamycin media) reprogrammed the transcriptome in aging cells (Fig. 5A and 5B). Above all, oxygenation broadly repressed lipid and purine biosynthesis transcripts (Fig. 5A), and in particular *OLE1*, a known hypoxia responder encoding the fatty acid desaturase (Kwast et al. 1999). Because lipid and purine metabolism are required for a long life (Arlia-Ciommo et al. 2018; Garay et al. 2014; Handee et al. 2016; Matecic et al. 2010), this repression may well contribute to the life span shortening of *WA-HPF1* cells. Overall, transcriptome changes implied that the life span shortening of WA-*HPF1* cells may arise due to reduced lipid and purine biosynthesis as well as oxidation of free or protein bound methionine.

## DISCUSSION

We found calorie restriction to delay chronological aging for each of the 1056 yeast genotypes studied. These overwhelmingly positive effect of calorie restriction is fully in line with reports on single gene knockouts, where calorie restriction extends chronological life span regardless of what gene is missing (Matecic et al. 2010). In contrast, calorie restriction only extends the replicative life span of approximately half of single gene knockouts, with many being negatively affected (Schleit et al. 2013). The diverging effects of calorie restriction on chronological and replicative life span may reflect the natural yeast life cycle; yeasts spend much time as starved, quiescent cells (Liti 2015) and survival in this state may be very strongly selected, but the aged, extensively replicated state is extremely rare and unlikely to be selected.

Chronological life span was extremely genotype dependent. Using genome-wide linkage analysis, we found a total of 30 distinct QTLs explaining up to 41% of life span variation. Among those, two major QTLs drove most of the life span variation (~30% and ~20%), while the remaining QTLs were more time and environment specific and contributed less (~3% each). We linked the two major QTLs to the cell wall encoding genes *FLO11* (chrIX) and *HPF1* (chrXV), with West African (WA) alleles having a pronounced and dominant life span shortening effect. Cell wall proteins have previously been linked to yeast life span through their effect on cell wall integrity. The cell wall integrity pathway reacts to cell wall perturbations, induced by e.g. heat shock or starvation (Krause and Gray 2002), and regulates both chronological and replicative life spans (Kaeberlein and Guarente 2002; Matecic et al. 2010; Ray et al. 2003; Stewart et al. 2007). We found that the life span shortening induced by the WA-*HPF1* was not by cell wall damage, but by a dramatic shift in life style. Cells carrying a North American (NA) *HPF1* had long, sedentary lives in sedimented yeast populations, while cells carrying a WA-*HPF1* lived shorter as buoyant, free-floating yeasts. The buoyancy was directly causing the shorter life span since enforcing a free-floating state of cells carrying the NA-*HPF1*, through vigorous shaking, shortened the life span to the same levels as that of cells carrying the WA-*HPF1*. We observed no strong buoyancy effect of *FLO11* variation although *FLO11* induced buoyancy has been previously reported (Fidalgo et al. 2006).

The reason why buoyancy shortens life span is likely by shifting cells from semi-anaerobic state within yeast sediments, to a highly oxygenated state while floating. Higher oxygen exposure means higher oxidative damage by reactive oxygen species, an oft-suggested explanation for life span shortening (Fabrizio et al. 2003; Longo et al. 1999). We found buoyant yeasts to broadly induce cellular systems dealing with methionine oxidation and depletion. Non-oxidized methionine is essential to the proper folding and function of proteins, synthesis of the central signalling molecule S-adenosine methionine and the maintenance of glutathione pools, a key redox buffer (Brown-Borg and Buffenstein 2017), helping to explain why buoyancy shortens life span. Rapamycin supplementation abolished the induction of the methionine oxidation response in buoyant yeasts, implying that rapamycin counters methionine oxidation and potentially explaining some of its life span extending effect. Buoyancy also broadly repressed lipid and purine biosynthesis genes. Both lipid and purine metabolisms have previously been linked to life span extension (Arlia-Ciommo et al. 2018; Matecic et al. 2010; Mohammad et al. 2018). Why this occurs in buoyant cells is unclear. It is possible that lipid and purine oxidation, caused by the excess load of reactive oxygen species in oxygen exposed buoyant yeasts, shortens life span by damaging membranes, signalling molecules and nucleotides. In addition, several studies connect methionine and lipid metabolisms, suggesting that early methionine perturbation during aerobic growth distorts lipid metabolism in aged cells (Hasek et al. 2013; Lee et al. 2014; Zhou et al. 2016).

*HPF1* and *FLO11*, like the vast majority of genes encoding cell wall proteins, contain intragenic tandem repeats (Verstrepen et al. 2005). Intragenic tandem repeats are dynamic in size, both due to strand-slippage during replication and ectopic recombination (Fan and Chu 2007; Pâques, Leung, and Haber 1998), and were found to fuel rapid yeast evolution (Gemayel et al. 2012; Verstrepen et al. 2005). Here, we showed that expansion of the N-terminal intragenic repeats within WA-*HPF1* was sufficient to shift yeasts towards a buoyant life style, which negatively impacted life span. The majority (14/21) of amino acids in the expanded N-terminal repeat motif are serines or threonines, a huge overrepresentation compared to the 12% expected by their general prevalence in proteins (Kozlowski 2016). Serine and threonine are unique among amino acids in containing hydroxyl groups that directly facilitate hydrogen bonding with surrounding water molecules, an effect known to enhance the solubility of organic particles. An enticing possibility is therefore that the serine/threonine richness of the *Hpf1p* intragenic repeat expansion directly induces buoyancy. Furthermore, serine and threonine residues are highly O-glycosylated in the cell wall and in secreted proteins. Repeat expansion could thus increase cell wall glycosylation, which is predicted to improve solubility. In this model, the dynamic repeat expansions and contractions of *HPF1* may serve as a life style switch, allowing rapid shifts between buoyant and sedentary life styles in evolution as dictated by fluctuating, opposing selection pressures. We note that the biochemical properties of the yeast cell wall have been linked to buoyancy (DeSousa et al. 2003; Fidalgo et al. 2006; Palmieri, Greenhalf, and Laluce 1996), albeit through hydrophobicity rather than by serine/threonine induced hydrogen bond formation with water molecules.

The *HPF1* induced shift to a buoyant life style with concomitant life span shortening due to methionine, lipid and purine oxidation has no immediate parallel in multicellular organisms. Nevertheless, intragenic tandem repeats are enriched in human genes encoding extracellular proteins (Gemayel et al. 2010; Legendre et al. 2007). Mammalian cells maintain a complex relationship with the surrounding extracellular matrix. For instance, stem cell fate and proliferation are strongly determined by physical constraints, cellular shape, and environmental cues, including oxygen availability, through interactions with the surrounding matrix (Campisi 2001; Guilak et al. 2009; Rando 2006). The crosslinking theory of aging postulates that the progressive crosslinking of the extracellular matrix impairs tissue homeostasis and drives aging (Bjorksten 1968). Notably, fibroblast senescence can be reversed by culturing old cells in a young extracellular matrix (Choi et al. 2011). It is not inconceivable that intragenic repeat expansions also in human extracellular proteins modulate cell-matrix interactions with profound effects on cellular life span.

In a broader perspective, our understanding of the biological role of intragenic tandem repeats is still rudimentary. The study of the genotype-phenotype map has largely been limited to SNPs, CNVs and aneuploidies. Nonetheless, there is an increasing awareness that intragenic tandem repeat polymorphisms generate important functional variability. In addition to human neurodegenerative diseases such as Huntington’s chorea or Fragile X syndrome (Gemayel et al. 2010; Orr and Zoghbi 2007), tandem repeat variations have been reported to drive the evolution of pathogenic bacteria (Stern et al. 1986), circadian clocks (Sawyer et al. 1997), and organismal morphology (Fondon and Garner 2004). Roughly, 17% of genes in the human genome contain intragenic tandem repeats (Legendre et al. 2007), and these are likely to have far-reaching effects on human biology. The ongoing development of long read sequencing will allow detecting intragenic repeat polymorphism and help uncover associations to many classes of phenotypic variation, as demonstrated in our study.

## MATERIALS AND METHODS

### Strains

Phased Outbred Lines (POLs) were derived from a cross between a North American (NA) oak tree strain (YPS128) and a West African (WA) palm wine strain (DBVPG6044) (Liti et al. 2009). Heterothallic ancestral parents carrying *LYS2* or *URA3* at the *LYS2* locus (Cubillos, Louis, and Liti 2009) were first mated to generate a pool of F1 hybrids. The pool was cycled through 12 rounds of alternating random mating, diploid selection, meiosis, sporulation, and haploid selection, resulting in a final pool of F12 outbred haploids (Parts et al. 2011). 86 F12 haploid segregants of each mating type were isolated, sequenced and their genotype was inferred using a set of 52466 markers (Illingworth et al. 2013). Phased Outbred Lines (POLs) were obtained as described (Hallin et al. 2016) with minor modifications. F12 segregants of opposite mating type were randomly paired and mated (YPD) in 96 well plates, and 1056 unique diploids with known, phased genomes were then selected during 3 consecutive diploid selective cultivations on minimal media. Diploids were arrayed, stored and analysed in 12×96 well plates, each plate containing 8 (3 each of NA/NA and WA/WA, and 2 of NA/WA) internal controls used for life span normalisation. Reciprocal hemizygotes at the *HPF1* and *FLO11* loci were constructed in a NA/WA diploid hybrid, using genetically tractable NA and WA haploids, as described (Cubillos, Louis, and Liti 2009). Native *HPF1* and *FLO11* genes were deleted in haploids by homologous recombination with a NatMX4 cassette, using the lithium acetate/PEG transformation protocol (Cubillos, Louis, and Liti 2009), before being mated to the appropriate counterpart to generate diploids hemi- or homozygote for *HPF1* and *FLO11*. *HPF1* allele swapping was performed in two steps. First, part of *HPF1* (N-terminal repeats, C-terminal repeats, or internal part (Fig. 3C) was deleted in NA and WA haploids using homologous recombination with a *URA3* cassette. Then, the *HPF1* segments to be swapped were PCR amplified from the desired alleles with Platinum Superfi (Invitrogen) DNA polymerase and swapped into the orthologous position of the recipient strain using homologous recombination (targeting identical non-repeated sequences for both alleles) and selected on 5-FOA 1 g/L. Strains obtained were then mated to *hpf1*::NatMX4 haploids to generate the indicated diploids. Complete deletion of *HPF1* in WA and NA diploids was performed in two consecutive transformation steps, replacing both copies with NatMX4 and KanMX4 cassettes. A summary of strains and primers used in this study can be found in tables S1 and S2, respectively.

### Media

YPD (1% yeast extract, 2% peptones, 2% dextrose, 2% agar (MP Biomedicals)) was used for all matings. Mated cells were streaked on synthetic minimal medium (2% dextrose, 0,675% yeast nitrogen bases (Formedium), pH set to 6.0 with 2.5M NaOH), to select for diploids. Life span was estimated in: i) calorie rich synthetic dextrose complete (SDC) media (2% dextrose, 0.675% yeast nitrogen base (Formedium), 0.088% complete amino acid supplement (Formedium), pH set to 6.0 with 2.5M NaOH), ii) calorie restricted (CR) media (SDC as above, but with 0,5% dextrose instead of 2%) (Jiang 2000; Lin, Defossez, and Guarente 2000; Smith et al. 2007), and iii) rapamycin (RM) supplemented media (SDC supplemented with 0,025 µg/mL rapamycin (Sigma-Aldrich)) (Li et al. 2019; Vázquez-García et al. 2017).

### Chronological lifespan assay

Cells cultivated overnight in calorie-rich (SDC) media were diluted (1:100) in 200 µL of either fresh SDC, or CR, or RM media in a 96 well plate. Cultivation plates were sealed with adhesive aluminium foil to prevent evaporation and incubated at 30°C. Aging was considered to start at saturation of the culture, 72 hours post inoculation (Fabrizio and Longo 2007), and cells were kept in saturated media for the whole duration of the experiment unless otherwise specified. When CLS was performed in water (Fig. S4A only), 72 hours stationary cultures were centrifuged and cells were washed 3x before being resuspended and kept in the same volume of distilled water.

Aging was measured as viable cells (%) by flow cytometry based on the uptake of the fluorescent molecules propidium iodide (PI) and YO-PRO-1 iodide (YP). Propidium iodide and YO-PRO-1 are membrane-impermeable nucleic acid binding molecules that enter into necrotic but not into alive cells. Therefore, non-fluorescent cells are alive, while fluorescent cells are not. YO-PRO-1 penetrates also into apoptotic cells (Herker et al. 2004; Wlodkowic, Skommer, and Darzynkiewicz 2009). At each aging time point (7, 21, 35 days after entry into quiescence unless otherwise stated), 5 µL of cells were transferred into 100 µL of staining solution (Phosphate Buffer Saline + 3 µM propidium iodide (Sigma) + 200nM YP (Invitrogen)) in a 96 well plate and incubated for 5 min in the dark at 30°C. The samples were analysed on a FACS-Calibur flow cytometer (Becton Dickinson) using a High Throughput Sampler (Becton Dickinson) device to process 96 well plates and detect fluorescence with FL-1 (YP) and FL-3 (PI) channels. POLs experiments, where each allele is present and therefore replicated in many lineages, were run in single replicate and viability estimates were normalized to those of 8 internal controls run on the same plate. All other experiments were run at least in triplicates.

### Linkage analysis

Linkage analysis was performed as described (Hallin et al. 2016). Briefly, mapping of life span QTLs was done using the normalized POL life spans, the R/qtl package in R (Broman et al. 2003), and the marker regression method in the scanone function. Significance thresholds were calculated with 1,000 permutations to call QTLs with a significance level of 0.05. Confidence intervals for the peaks were calculated using a 1.8-LOD drop using the lodint function in R/qtl. We corrected for population structure by using the deviation of the lifespan of each POL from the parent mean.

### RNAseq

We extracted RNA from SDC and RM cultivated cells in exponential phase and after 7 days of aging using the Kapa Biosystems hyperplus kit with Riboerase, as per user’s instructions at 500nG input. RNA integrity was assessed on an Agilent Bioanalyzer using the RNA pico kit to determine RIN scores. Library quality was assessed using Agilent Bioanalyzer using the DNA 1000 Kit. Libraries were quantified using Kapa Biosystems library quantification kit. The libraries were normalized, pooled and loaded onto a NextSeq 500/550 High Output v2 kit (300 cycles) flow cell and sequenced on a Next seq 500 instrument. We performed transcript-level abundance quantification by pseudo-aligning RNA-seq reads to the coding sequences of SGD yeast reference gene set, using Kallisto (v0.44.0) (Bray et al. 2016). In this way, we obtained the Transcripts Per Kilobase Million (TPM) value for each gene in each sample as its normalized expression level, which is directly comparable both among different genes and among different samples. Sleuth (0.30.0) (Pimentel et al. 2017) was further used to assess the statistical difference of the same gene between different samples by decoupling biological differences from experimental noise. False discovery rate (FDR; α = 0.05) adjustment (Benjamini and Hochberg 1995) was further applied for multiple test correction.

Total RNA was extracted from two biological replicates either during exponential growth or after 7 days of aging in SDC or in RM. Global expression level was analysed by pairwise comparison of the NA/WA hybrids hemizygotes for WA and NA-*HPF1* within each environment and time point to identify differentially expressed transcripts.

Standard GO term analysis was performed on >2-fold differently expressed genes with the GO Term Finder tool available at SGD, with an FDR corrected α threshold of 0.01.

Raw RNAseq reads are available at the NCBI Sequence Read Archive under accession PRJNA544860.

## Acknowledgments

This work was supported by Agence Nationale de la Recherche (ANR-11-LABX-0028-01, ANR-13-BSV6-0006-01, ANR-15-IDEX-01, ANR-16-CE12-0019 and ANR-18-CE12-0004) and the Swedish Research Council (2014-6547, 2014-4605 and 2018-03638).

