## Supplemental Figures for "Intragenic repeat expansions control yeast chronological aging"

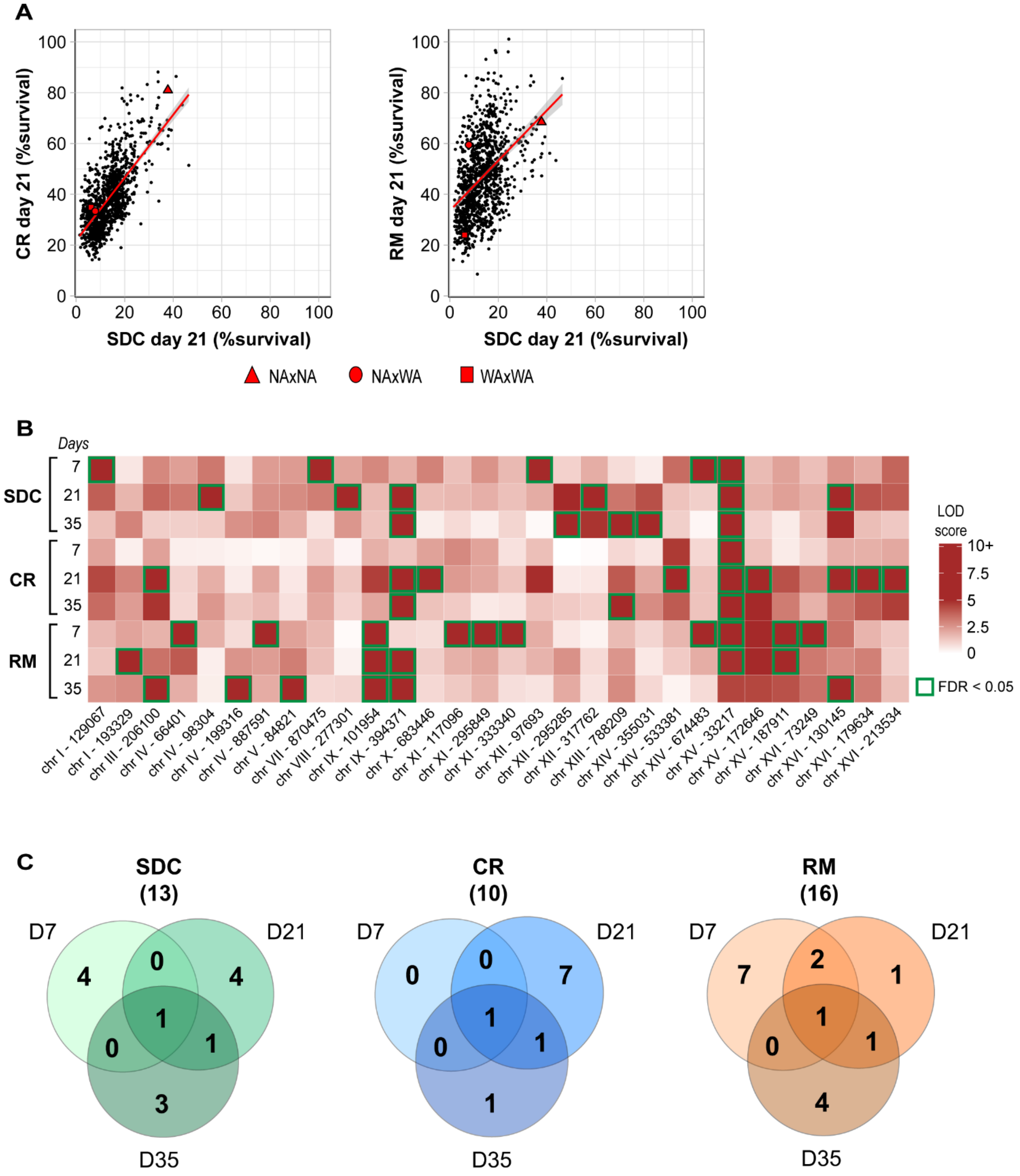


**Fig S1. Genetic, environmental and time variation in chronological life span**

**Figure S1. Genetic, environmental and time variation in chronological life span**

**(A)** Scatter plots of CLS of selected environments and time points for the 1056 segregants (reported in Fig. 1A) underlie CLS correlation across conditions. Parental strains and F1 hybrid are indicated with red symbols. Linear regression is represented as a red line with 95% confidence interval. **(B)** Summary heat map of the 30 distinct QTL regions that were mapped. Green frames indicate conditions for which QTLs were significant (FDR < 0.05). **(C)** Venn diagram summarizing the distribution of QTLs across time points for each environment. D7, D21, and D35 refers to 7, 21, and 35 days of chronological aging, respectively. The value in parenthesis indicates the total number of QTLs found per environment.

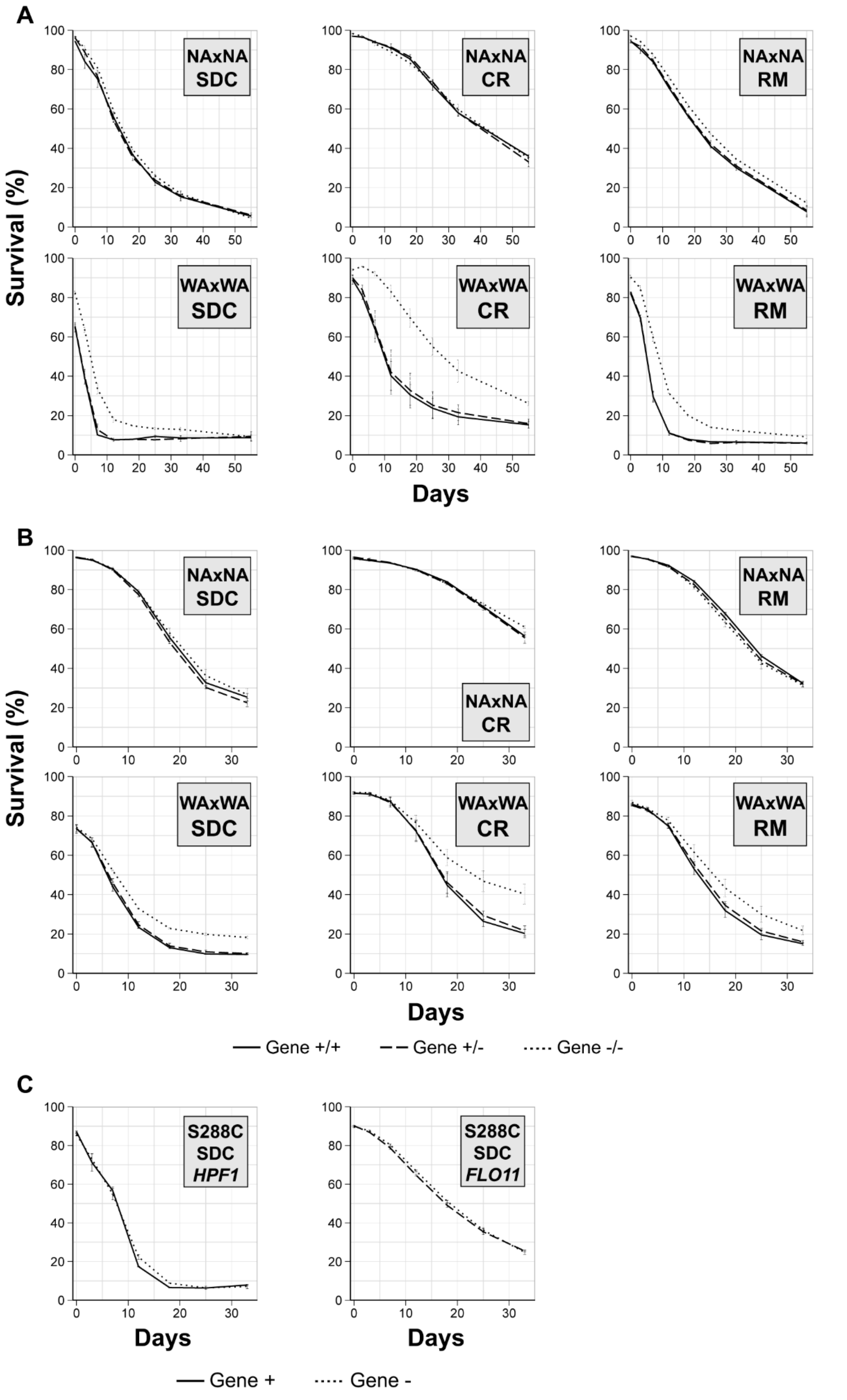


**Fig S2. Effect of *HPF1* and *FLO11* deletions on homozygote parental strains and lab strain S288C**

**Figure S2. Effect of *HPF1* and *FLO11* deletions on homozygote parental strains and lab strain S288C**
**(A)** Effect of *HPF1* deletion on NA and WA homozygote parents. Strain background and environmental condition are indicated in grey boxes. Solid lines, wild-type; dashed lines, one copy deleted; dotted lines, complete deletion. **(B)** Effect of *FLO11* deletion on NA and WA homozygote parents (see above). **(C)** Effect of *HPF1* and *FLO11* deletion on the lab strain S288C in SDC. Gene deleted is indicated in the grey box. Solid lines, wild-type; dotted lines, complete deletion.

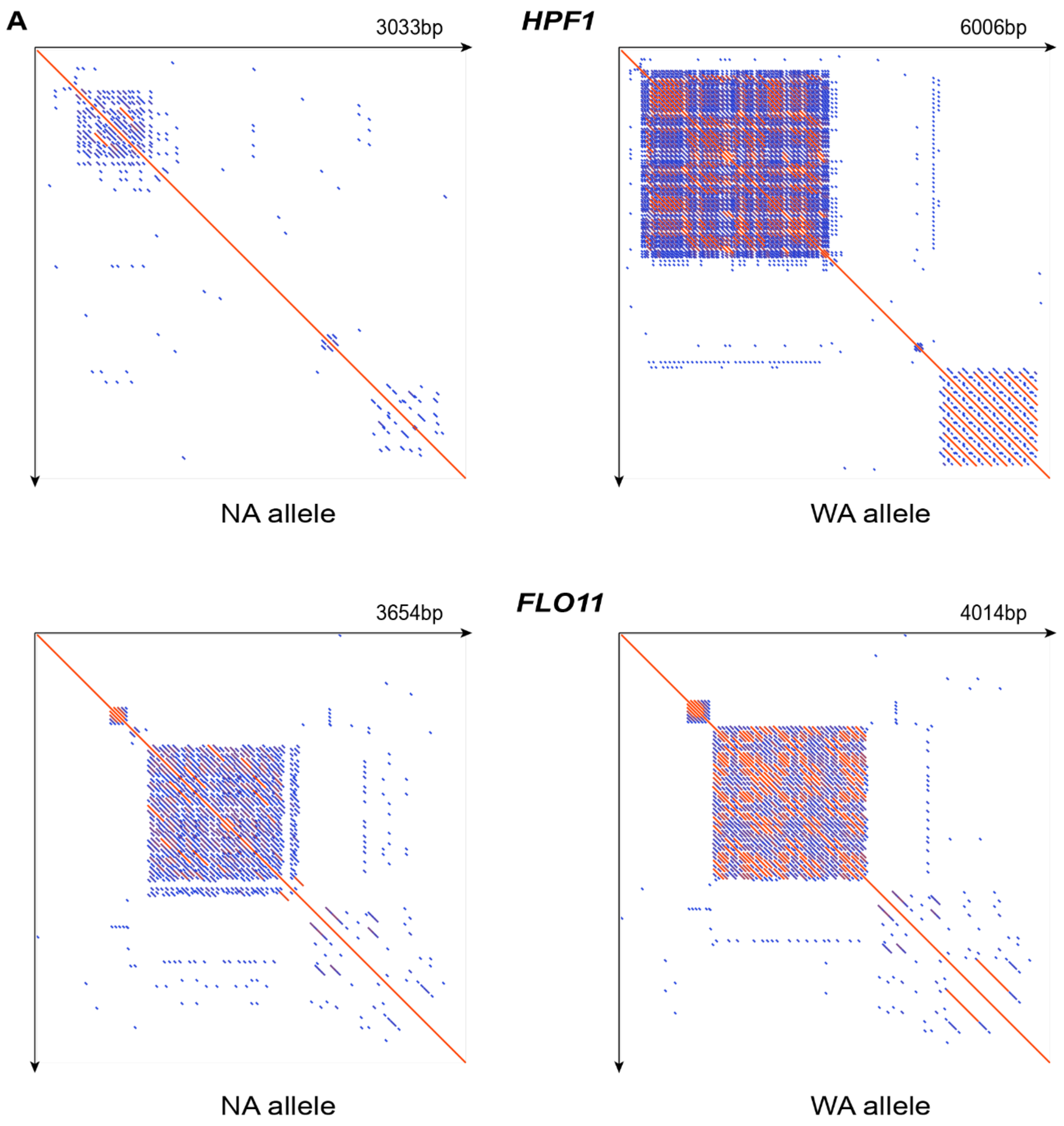

A
HPF1
3033bp
NA allele
6006bp
WA allele
FLO11
3654bp
NA allele
4014bp
WA allele

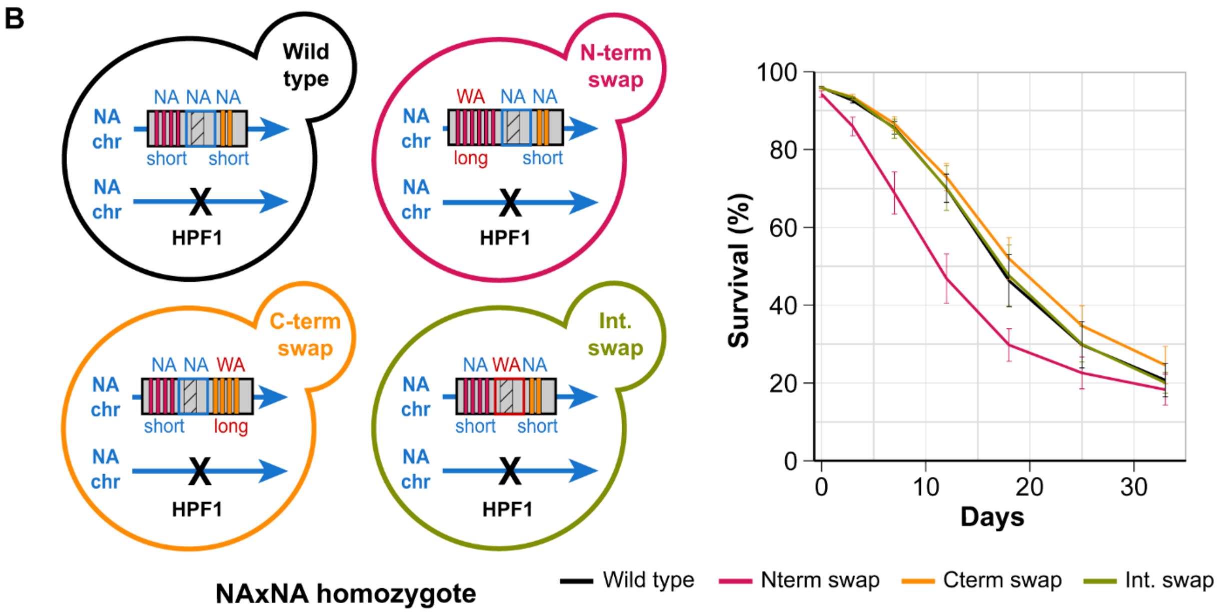


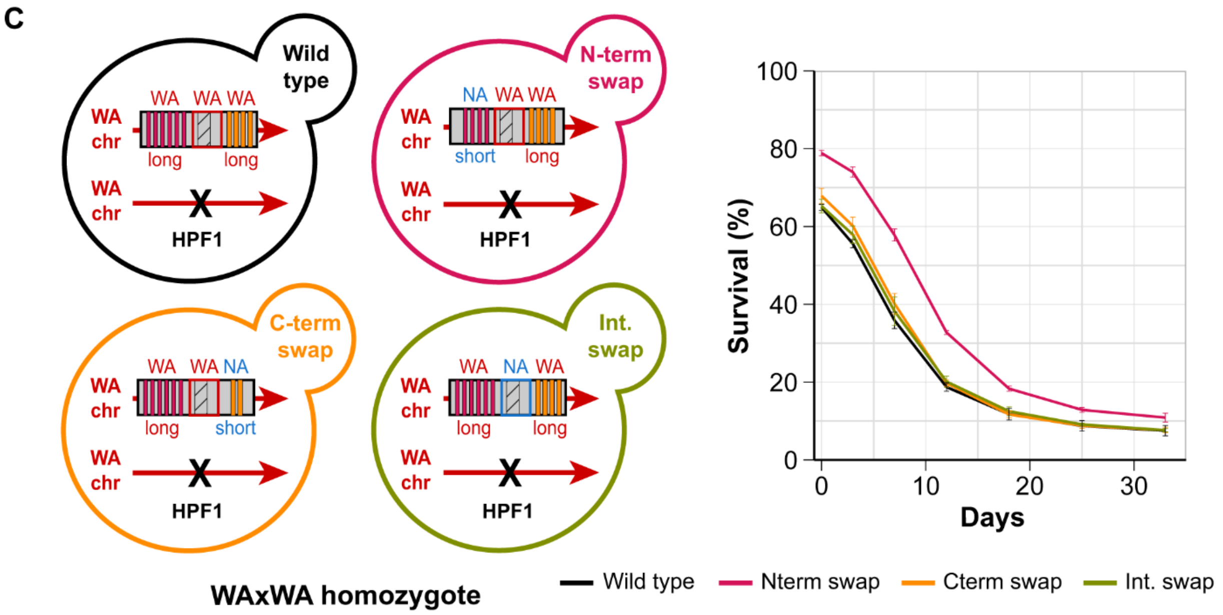


**Fig S3. N-terminal repeat expansion within WA-*HPF1* shortens chronological life span**

**Figure S3. N-terminal repeat expansion within WA-*HPF1* shortens chronological life span**

**(A)** Self alignment of *HPF1* (top) and *FLO11* (bottom) ORFs for the NA (left) or WA alleles (right). Created with the software Geneious. **(B)** Allele swapping of different *HPF1* parts effect on CLS. The NA/NA homozygote was engineered as described in figure 3C. The various constructions are schematically represented on the left. **(C)** Similar to panel B for the WA/WA homozygote.

A

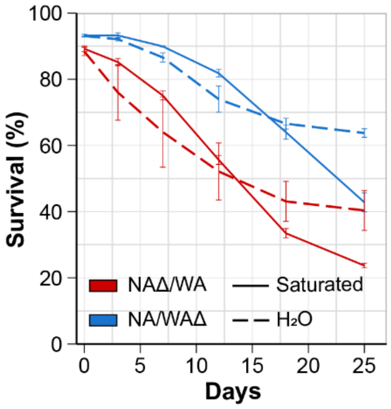


B

| Timepoint | SDC | | CR | | RM | |
|---|---|---|---|---|---|---|
| | NAΔ/WA | NA/WAΔ | NAΔ/WA | NA/WAΔ | NAΔ/WA | NA/WAΔ |
| Inoculation | 6.00 | 6.00 | 6.29 | 6.29 | 6.00 | 6.00 |
| Mid-Exponential | 3.98 | 4.10 | 4.54 | 4.71 | 3.97 | 4.04 |
| 7 days | 3.52 | 3.53 | 4.69 | 4.68 | 3.47 | 3.50 |
| 21 days | 3.50 | 3.51 | 4.63 | 4.88 | 3.44 | 3.41 |

**Fig S4. Buoyancy shortens life span without altering the extracellular medium composition or acidity**

**Figure S4. Buoyancy shortens life span without altering the extracellular medium composition or acidity**
**(A)** CLS of *HPF1* reciprocal hemizygote strains in water (Δ indicates which *HPF1* allele was deleted). Cells were pre-grown 3 days in SDC before washed and resuspended in water or kept in saturated media. Error bars represent standard deviations. **(B)** pH of *HPF1* reciprocal hemizygote cultures during CLS. Cells were grown and aged either in SDC, CR, or RM, as indicated. Inoculation refers to the pH at the time of initial incubation, mid-exponential corresponds to 7 hours post-inoculation.
