## Supplemental Table 6 for "Intragenic repeat expansions control yeast chronological aging"

**TABLE S6 – *FLO11* and *HPF1* DNA sequences (ORF)**

**>*FLO11*_DBVPG6044**

ATGCAAAGACCATTTCTACTCGCTTATTTGGTCCTTTCGCTTCTATTTAACTCAGCTTTGGGTTTTCCAACTGCACTAGTTCCTAGATGCTCCGAAGGAACTAGCTGTAATTCTATCGTTAATGGCTGTCCCAACTTAGACTTCAATTGGCACATGGACCAACAGAACATCATGGAGTATACTTTGGATGTGACTTCTGTTTCTTGGGTTCAAGACAACACGTACCAAATCACTGTTCATGTCAAAGGTAAAGAAAACATTGACCTAAAATATCTATGGTCTTTGAAAATCATTGGTGTTACTGGTCCAAAAGGTACCGTCCAACTATACGGTTACAACGAAAATACCTATTTGATTGACAACCCAACTGATTTCACAGCCACTTTTGAAGTCTATGCCACACAAGATGTCAACAGCTGTCAGGTGTGGATGCCTAACTTCCAAATTCAATTCGAGTATTTGCAAGGTAGTGCCGCTCAATATGCAAGCTCTTGGAAATGGGGAACTACATCTTTTGATTTGTCTACTGGTTGTAACAACTATGACAATCAAGGCCACTCTCAAACAGATTTCCCAGGCTTCTATTGGAACATAGATTGTGACAACAATTGTGGCGGTACGAAGTCATCTACCACTACATCAACTAGTACTTCCGAGTCATCTACCACTACATCAACTAGTACTTCCGAGTCATCTACCACTACATCAACTAGTACTTCCGAGTCATCTACCACTACATCAACTAGTACTTCCGAGTCATCTACCACTACATCTAGCACTTCCGAGTCATCTACCACTACATCAACTACCACTTCAGAGTCATCTACATCATCATCAACCACCGCTCCTGCTACACCAACCACTACCGAAAGCTCTTCTGCTCCAGTACCAACTCCATCAAGCTCTACTACTGAAAGCTCTTCTGCTCCAGTAACCAGCTCCACCACTGAAAGCTCTTCTGCTCCAGTAACCAGCTCCACCACTGAAAGCTCTTCTGCTCCAGCTCCAACTCCATCCAGCTCTACTACCGAAAGCTCTTCTGCTCCAGTATCCAGCTCCACCACTGAAAGCTCTTCTGCTCCAGCTCCAACTCCATCCAGCTCTACTACCGAAAGCTCTTCTGCTCCAGTAACCAGCTCCACCACTGAAAGCTCTTCTGCTCCAGTAACCAGCTCCACCACTGAAAGCTCTTCTGCTCCAGTAACCAGCTCCACCACTGAAAGCTCTTCTGCTCCAGTACCAACTCCATCAAGCTCTACTACTGAAAGCTCTTCTGCTCCAGTAACCAGCTCCACCACTGAAAGCTCTTCTGCTCCAGTACCAACTCCATCAAGCTCTACTACTGAAAGCTCTTCTGCTCCAGTACCAACTCCATCAAGCTCTACTACTGAAAGCTCTTCTGCTCCAGTACCAACTCCATCCAGCTCTACTACCGAAAGCTCTTCTGCTCCAGTACCAACTCCATCAAGCTCTACTACTGAAAGCTCTTCTGCTCCAGTAACCAGCTCCACCACTGAAAGCTCTTCTGCTCCAGTAACCAGCTCCACCACTGAAAGCTCTTCTGCTCCAGCTCCAACTCCATCCAGCTCTACTACCGAAAGCTCTTCTGCTCCAGTATCCAGCTCCACCACTGAAAGCTCTTCTGCTCCAGCTCCAACTCCATCCAGCTCTACTACCGAAAGCTCTTCTGCTCCAGTAACCAGCTCCACCACTGAAAGCTCTTCTGCTCCAGTACCAACTCCATCAAGCTCTACTACTGAAAGCTCTTCTGCTCCAGTAACCAGCTCTACTACTGAAAGCTCTTCTGCTCCAGTACCAACTCCATCAAGCTCTACTACTGAAAGCTCTTCTGCTCCAGTAACCAGTTCCACCACTGAAAGCTCTTCTGCTCCAGTACCAACTCCATCCAGCTCTACCACTGAAAGCTCTTCTGCTCCAGCTCCAACTCCATCCAGCTCTACTACCGAAAGCTCTTCTGCTCCAGTATCCAGCTCCACCACTGAAAGCTCTTCTGCTCCAGCTCCAACTCCATCCAGCTCTACTACCGAAAGCTCTTCTGCTCCAGTAACCAGCTCCACCACTGAAAGCTCTTCTGCTCCAGTAACCAGCTCCACCACTGAAAGCTCTTCTGCTCCAGTACCAACTCCATCAAGCTCTACTACTGAAAGCTCTTCTGCTCCAGTAACCAGCTCCACCACTGAAAGCTCTTCTGCTCCAGTACCAACTCCATCAAGCTCTACTACTGAAAGCTCTTCTGCTCCAGTAACCAGCTCCACCACTGAAAGCTCTGTAGCACCAGTACCAACCCCATCTTCCTCTAGCAACATCACTTCCTCCGCTCCATCATCATCCAAATACCCTGGCAGTCAAACAGAAACCTCTGTTTCTTCTACAACCGAAACTACCATTGTTCCAACTACAACTACGACTTCTGTCACTACACCATCAACAACCACTATTACCACTACGGTTTGCTCTACAGGAACAAACTCTGCCGGTGAAACAACCTCTGGATGCTCTCCAAAGACCGTTACAACTACTGTTCCAACTACAACTACGACTTCTGTCACTACATCATCAACAACCACTATTACTACTACGGTTTGCTCTACAGGAACAAACTCTGCCGGTGAAACTACTTCTGGATGCTCTCCAAAGACCATTACAACTACTGTTCCATGTTCAACCAGTCCAAGCGAAACCGCCTCGGAATCAACAACCACTTCACCTACCACACCTGTAACTACAGTTGTCTCAACCACCGTCGTTACTACTGAGTATTCTACTAGTACAAAACCAGGTGGTGAAATTACAACTACATTTGTCACCAAAAACATTCCAACCACTTACCTAACCACAATTGCTCCAACTCCATCAGTCACTACGGTTACCAATTTCACCCCAACCACTATTACTACTACGGTTTGCTCTACAGGTACAAACTCTGCCGGTGAAACTACCTCTGGATGCTCTCCAAAGACTGTCACAACCACTGTTCCTTGTTCAACTGGTACTGGCGAATACACTACTGAAGCTACCACCCCTGTTACAACAGCTGTCACAACCACCGTTGTTACCACTGAATCATCTACGGGTACTAACTCCGCTGGTGAGACGACAACTGGTTACACAACAAAGTCTGTACCAACCACCTATGTAACCACTTTGGCTCCAAGTGCACCAGTAACTCCTGCCACTAATGCCGTACCAACTACAATAACCACTACTGAATGTTCTGCTGCTACAAACGCTGCCGGTGAAACTACATCTGTATGCTCTGCTAAGACTATCGTAAGTTCTGCAAGCGCAGGCGAAAACACCACCCCTGTCACGACAGCTGTCACAACCACCGTTGTTACCACTGAATCATCTACGGGTACTAACTCCGCTGGTGAGACGACAACTGGTTACACAACAAAGTCTGTACCAACCACCTATGTAACCACTTTGGCTCCAAGTGCACCAGTAACTCCTGCCACTAATGCCGTACCAACTACAATAACCACTACTGAATGTTCTGCTGCTACAAACGCTGCCGGTGAAACTACATCTGTATGCTCTGCTAAGACTATCGTAAGTTCTGCAAGCGCAGGCGAAAACACCACCCCTGTCACGACAGCTATTCCAACCACAGTTGTTACCACTGAGTCATCTGTTGGTACTAACTCCGCTGGCGAAACAACAACTGGTTACACAACCAAGTCCATCCCAACCACTTACATAACCACTTTGATTCCAGGTTCAAATGGTGCCAAGAATTACGAAACTGTGGCCACAGCAACCAACCCTATTTCAATCAAGACTACATCCCAACTAGCTACAACAGCTTCTGCTTCTAGCATGGCTCCCGTTGTCACATCTCCATCTCTAACTGGTCCACTACAATCTGCTTCTGGTTCTGCAGTCGCTACATACTCTGTTCCTTCTATCTCGAGTACTTACCAAGGTGCTGCTAATATCAAGGTTCTTGGAAACTTTATGTGGTTGCTACTCGCTCTTCCAGTTGTATTCTAA

**>*FLO11*_YPS128**

ATGCAAAGACCATTTCTACTCGCTTATTTGGTCCTTTCGCTTCTATTTAACTCAGCTTTGGGTTTTCCAACTGCACTAGTTCCTAGAGGATCCTCCGAAGGAACTAGCTGTAATTCTATCGTTAATGGCTGTCCCAACTTAGACTTCAATTGGCACATGGACCAACAAAATATCATGCAGTATACTTTGGATGTGACTTCCGTTTCTTGGGTTCAAGACAACACATACCAAATCACTATTCATGTCAAAGGTAAAGAAAACATTGACCTAAAATATCTATGGTCTTTGAAAATCATTGGTGTCACTGGTCCAAAAGGTACCGTCCAACTATACGGTTACAACGAAAATACCTATTTGATTGACAACCCAACTGATTTCACAGCCACTTTTGAAGTCTATGCCACACAAGATGTCAACAGCTGTCAGGTGTGGATGCCTAACTTCCAAATTCAATTCGAGTATTTGCAAGGTAGTGCCGCTCAATATGCAAGCTCTTGGAAATGGGGAACTACATCTTTTGATTTGTCTACTGGTTGTAACAACTATGACAATCAAGGCCACTCTCAAACAGATTTCCCAGGCTTCTATTGGAACATAGATTGTGACAACAATTGTGGCGGTACGAAGTCATCTACCACTACATCAACTAGTACTTCCGAGTCATCTACCACTACATCAACTAGTACTTCCGAGTCATCTACCACTACATCAACTAGTACTTCCGAGTCATCTACCACTACATCAACTACCACTTCAGAGTCATCTACATCATCATCAACCACCGCTCCTGCTACACCAACCACTACCTCATGCACTAAGGAAAAGCCTACACCCCCAACCACTACCTCATGCACAAAGGAAAAGCCTACACCTCCTCATCACGACACCACTCCATGTACAAAGAAGAAAACCACCACATCTAAGACATGCACTAAGAAGACTACTACTCCAGTACCAACCCCATCAAGCTCTACTACTGAAAGCTCTTCTGCTCCAGTACCAACCCCATCAAGCTCTACCACTGAAAGCTCTTCTGCTCCAGTAACCAGCTCTACTACCGAAAGCTCTTCTGCTCCAGTACCAACTCCATCAAGCTCTACCACTGAAAGCTCTTCTGCTCCAGTACCAACTCCATCCAGCTCTACTACTGAAAGCTCTTCTGCTCCAGTACCAACTCCATCAAGCTCTACTACTGAAAGCTCCTCTGCTCCAGCTCCAACTCCATCCAGCTCCACTACTGAAAGCTCCTCTGCTCCAGTATCCAGCTCTACTACTGAAAGCTCTTCTGCTCCAGTACCAACTCCATCAAGCTCTACTACTGAAAGCTCCTCTGCTCCAGCTCCAACTCCATCCAGCTCCACTACTGAAAGCTCCTCTGCTCCAGTATCCAGCTCTACTACTGAAAGCTCTTCTGCTCCAGTACCAACTCCATCCAGCTCTACCACTGAAAGCTCTTCTGTTCCAGTACCAACCCCATCAAGCTCTACTACTGAAAGCTCTTCTGCTCCAGTACCAACCCCATCAAGCTCTACCACTGAAAGCTCTTCTGCTCCAGTAACCAGCTCTACTACCGAAAGCTCTTCTGCTCCAGCTCCAACTCCATCCAGCTCTACTACTGAAAGCTCTTCTGCTCCAGCTCCAACTCCATCCAGCTCTACTACTGAAAGCTCTTCTGCTCCAGTAACCAGCTCTACCACTGAAAGCTCTTCTGCTCCAGCTCCAACTCCATCCAGCTCCACCACTGAAAGCTCTTCTGCTCCAGTACCAACTCCATCAAGCTCTACCACTGAAAGCTCTTCTGCTCCAGTACCAACTCCATCAAGCTCCACTACTGAAAGCTCTTCTGCTCCAGCTCCAACTCCATCCAGCTCCACTACTGAAAGCTCCTCTGCTCCAGTATCCAGCTCTACTACTGAAAGCTCTTCTGCTCCAGTACCAACTCCATCCAGCTCTACCACTGAAAGCTCCTCTGCTCCAGTAACCAGCTCTACTACCGAAAGCTCTTCTGCTCCAGCTCCAACTCCATCAAGCTCTACTACTGAAAGCTCCTCTGCTCCAGTATCCAGCTCTACTACTGAAAGCTCTGTAGCACCAGTACCAACCCCATCTTCCTCTAGCAACATCACTTCCTCCGCTCCATCTTCAACTCCATTCAGCTCTAGCACTGAAAGCTCTTCTGTTCCAGTATCCAGCTCCACCACTGAAAGCTCTGTAGCACCAGTACCAACCCCATCTTCCTCTAGCAACATCACTTCCTCCGCTCCATCATCATCCAAATACCCTGGCAGTCAAACAGAAACCTCTGTTTCTTCTACAACCGAAACTACCATTGTTCCAACTACAACTACGACTTCTGTCACTACACCATCAACAACCACTATTACCACTACGGTTTGCTCTACAGGAACAAACTCTGCCGGTGAAACAACCTCTGGATGCTCTCCAAAGACCGTTACAACTACTGTTCCAACTACAACTACGACTTCTGTCACTACATCATCAACAACCACTATTACTACTACGGTTTGCTCTACAGGAACAAACTCTGCCGGTGAAACTACTTCTGGATGCTCTCCAAAGACCATTACAACTACTGTTCCATGTTCAACCAGTCCAAGCGAAACCGCCTCGGAATCAACAACCACTTCACCTACCACACCTGTAACTACAGTTGTCTCAACCACCGTCGTTACTACTGAGTATTCTACTAGTACAAAACCAGGTGGTGAAATTACAACTACATTTGTCACCAAAAACATTCCAACCACTTACCTAACCACAATTGCTCCAACTCCATCAGTCACTACGGTTACCAATTTCACCCCAACCACTATTACTACTACGGTTTGCTCTACAGGTACAAACTCTGCCGGTGAAACTACCTCTGGATGCTCTCCAAAGACTGTCACAACCACTGTTCCTTGTTCAACTGGTACTGGCGAATACACTACTGAAGCTACCACCCCTGTTACAACAGCTGTCACAACCACCGTTGTTACCACTGAATCCTCTACGGGTACTAACTCCGCTGGTGAGACGACAACTGGTTACACAACAAAGTCTGTACCAACCACCTATGTAACCACTTTGGCTCCAAGTGCACCAGTAACTCCTGCCACTAATGCCGTACCAACTACAATAACCACTACTGAATGTTCTGCTGCTACAAACGCTGCCGGTGAAACTACATCTGTATGCTCTGCTAAGACTATCGTAAGTTCTGCAAGCGCAGGCGAAAACACCACCCCTGTCACGACAGCTATTCCAACCACAGTTGTTACCACTGAGTCATCTGTTGGTACTAACTCCGCTGGCGAAACAACAACTGGTTACACAACCAAGTCCATCCCAACCACTTACATAACCACTTTGATTCCAGGTTCAAATGGTGCCAAGAATTACGAAACTGTGGCCACAGCAACCAACCCTATTTCAATCAAGACTACATCCCAACTAGCTACAACAGCTTCTGCTTCTAGCATGGCTCCCGTTGTCACATCTCCATCTCTAACTGGTCCACTACAATCTGCTTCTGGTTCTGCAGTCGCTACATACTCTGTTCCTTCTATCTCGAGTACTTACCAAGGTGCTGCTAATATCAAGGTTCTTGGAAACTTTATGTGGTTGCTACTCGCTCTTCCAGTTGTATTCTAA

**>*HPF1*_DBVPG6044**

ATGGTCAAACCCATTGCTACACTTCAAGCCGTTTTGGCTTCGCTCCTTTACTCCCAAAGTAAATTGGGCCAATATTATACCCACAGTTCCTCAATCGCTAGTCACAGCTCCACTGCCGTTTCGTCAACTTCATCAGGTTCTGTTTCCATCAGTAGTTCTATTGTTGAGTCGACCTCATCTGCTTCTGATGTCTCGAGCTCTCTCACTGAGTTAACATCATCCTCCACCGAAGTCTCGAGCACCATTGCTCCATCAACCTCGTCCTCTGAAGTCTCGAGCTCTATTACTTCATCAGGCTCATCAGTCTCCGGCTCATCTTCTATTACTTCATCAGGCTCATCAGTCTCCAGTTCATCTTCTGTCACAGAATCAGGCTCATCCGCCCCAGGTTCATCTACTTCCATTACATCAGGTTCATCCTCCGCCACTGAATCAGGCTCATCAGTCTCCGGTTCATCTACTTCCATTACATCAGGCTCATCCTCCGCCACTGAATCAGGCTCATCAGTCTCTGGTTCATCTTCTGCCACAGAATCAGGCTCATCAGTCTCCGGTTCATCTACTTCCATTACATCAGGCTCATCCTCCGCCACTGAATCAGGCTCATCAGTCTCCGGTTCATCTACTTCCATTACATCAGGCTCATCCTCCGCCACTGAATCAGGCTCATCAGTCTCCGGTTCATCTACTTCCATTACATCAGGCTCATCCTCCGCCACTGAATCAGGCTCATCAGTCTCCGGTTCATCTACTTCCATTACATCAGGCTCATCCTCCGCCACTGAATCAGGCTCATCAGTCTCCGGTTCATCTACTTCCATTGCATCAGGCTCATCCTCCGCCACTGAATCAGGCTCATCAGTCTCCGGTTCATCTACTTCCATTACATTAGGCTCATCTTCTGTCACAGAATCAGGCTCATCAGTCTCCGGTTCATCTACTTCCATTACATCAGGCTCATCTTCTGTCACAGAATCAGGCTCATCCGCCCCAGGTTCATCTACTTCCATTACATCAGGCTCATCCTCTGCCACAGAATCAGGCTCATCCGCCCCAGGTTCATCTACTTCCATTACATCAGGTTCAACTTCTGTCATAGAATCAGGCTCATCAGTCTCCGGTTCATCTACTTCCATTACATCAGGCTCATCTTCTGTCACAGAATCAGGCTCATCCGCCCCAGGTTCATCTACTTCCATTACATCAGGCTCATCCTCTGCCACAGAATCAGGCTCATCCGCCCCAGGTTCATCTACTTCCATTACATCAGGCTCATCTTCTGTCACAGAATCAGGCTCATCCGCCCCAGGTTCATCTACTTCCATTACATCAGGCTCATCCTCTGCCACTGAATCAGGCTCATCCGCCCCAGGTTCATCTACTTCCATTACATCAGGTTCATCCTCCGCCACTGAATCAGGCTCATCCGCCCCAGGTTCATCTACTTCCATTACATCAGGTTCAACTTCTGCCACAGAATCAGGCTCATCCGCCCCAGGTTCATCTACTTCCATTACATCAGGTTCAACTTCTGCCACAGAATCAGGCTCATCCGCCTCCGGTTCATCCTCCGCCACAGAATCAGGCTCATCCGCCCCAGGTTCATCTACTTCCATTACATCAGGCTCATCTTCTGTCACAGAATCAGGCTCATCAGTCTCCGGTTCATCTACTTCCATTACATTAGGCTCATCTTCTGTCACAGAATCAGGCTCATCAGTCTCCGGTTCATCTACTTCCATTACATCAGGCTCATCTTCTGTCACAGAATCAGGCTCATCCGCCCCAGGTTCATCTACTTCCATTACATCAGGCTCATCCTCTGCCACAGAATCAGGCTCATCCGCCCCAGGTTCATCTACTTCCATTACATCAGGTTCAACTTCTGTCATAGAATCAGGCTCATCAGTCTCCGGTTCATCTACTTCCATTACATCAGGCTCATCTTCTGTCACAGAATCAGGCTCATCCGCCCCAGGTTCATCTACTTCCATTACATCAGGCTCATCCTCCGCCACAGAATCAGGCTCATCAGTCTCTGGTTCATCTTCTGCCACAGAATCAGGCTCATCAGTCTCCGGTTCATCTACTTCCATTACATCAGGCTCATCCTCCGCCACTGAATCAGGCTCATCAGTCTCCGGTTCATCTACTTCCATTACATCAGGCTCATCCTCCGCCACTGAATCAGGCTCATCAGTCTCCGGTTCATCTACTTCCATTACATCAGGCTCATCTTCTGCCACAGAATCAGGCTCATCAGTCTCCGGTTCATCTACTTCCATTACATCAGGTTCAACTTCTGCCACAGAATCAGGCTCATCCGCCTCCGGTTCATCTACTTCCATTACATTAGGCTCATCTTCTGTCACAGAATCAGGCTCATCAGTCTCCGGTTCATCTACTTCCATTACATCAGGCTCATCTTCTGTCACAGAATCAGGCTCATCAGTCTCCGGTTCATCTACTTCCATTACATCAGGCTCATCTTCTGTCACAGAATCAGGCTCATCAGTCTCCGGTTCATCTACTTCCATTACATTAGGCTCATCTTCTGTCACAGAATCAGGCTCATCAGTCTCCGGTTCATCTACTTCCATTACATCAGGCTCATCTTCTGTCACAGAATCAGGCTCATCCGCCCCAGGTTCATCTACTTCCATTACATCAGGCTCATCTTCTGTCACAGAATCAGGCTCATCCGCCCCAGGTTCATCTACTTCCATTACATCAGGCTCAACTTCTGCCACTGAATCAGGCTCATCCGCCCCAGGTTCATCTACTTCCATTACATCAGGTTCAACTTCTGCCACAGAATCAGGCTCATCCGCCTCCGGTTCATCTTCTGCCACAGAATCAGGCTCATCCGCCTCCGGTTCATCTTCTGCCACAGAATCAGGCTCATCCTCATCAGCATCTGAATCATCTATCACACAATCTGGTACCGCTTCCGGTTCATCAGCCTCCAGCACGTCCGGTTCTGTTACACAATCTGGTTCCTCCGTTTCCGGTTCATCAGCTTCTTCTGCTCCAGGTATCTCGAGTTCAATTCCTCAATCAACCTCATCGGCTTCCACTGCCTCCGGTTCTATCACCTCCGGTACCTTAAGTTCTATTACCTCTTCGGCTTCTAGTGCAACTGCAACTACTTCCAACTCTCTTTCTTCCAGCGACGGTACCGTTTACTTGCCATCCACAACAATTAGCGGTGATCTCAGAGTTACTGGTAAAGTAATTGCAACCGAGCCCGTGGAAGTCGCTGCCGGTGGTAAGTTGACTTTACTTGACGGTGAAAAATACGTCTTCTCATCTGATCTAAAAGTCTACGGTGACTTGCTTGTGAAAAAGTCCAAAGAAACCTATCCAGGTACCGAATTCGACATCTCCGGTGAAAACTTTGACGTGACCGGTAACTTCAACGCTGAAGAATCCGCTGCCACCTCTGCATCCATCTACTCCTTCACTCCAAGTTCTTTTGACAACAGTGGTGACATTTCCTTAAGTCTATCAAAGTCCAAGAAGGGTGAAGTCACTTTCTCTCCATACTCCAATTCTGGTGCCTTCTCTTTCTCGAACGCCATTCTCAACGGTGGTTCTGTTTCCGGTCTACAACGTAGAGACGACACTGAAGGTTCAGTAAACAACGGTGAAATTAACCTAGACAATGGAAGTACCTACGTCGTTGTCGAACCAGTTTCTGGAAGTGGTACAATCAACATCATCTCTGGCAACCTTTACTTGCACTATCCAGACACCTTTACTGGCCAAACTGTTGTATTCAAGGGTGAAGGTGTTCTTGCCGTTGACCCTACCGAAAGCAACACTACTCCTATCCCTGTGGTTGGATACACTGGTGAAAACCAAATCGCCATTACAGCAGATGTAACTGCTCTTTCTTACGACAGTGCTACTGGTGTTTTAACTGCAACACAAGGCAACTCACAATTCTCCTTCTCTATTGGTACTGGATTCTCCAGTTCTGGTTTCAACGTCTCCGAAGGAACATTTGCTGGTGCCTATGCTTATTATCTAAATTACGGAGGTGTTGTTGCTTCCAGCGCTACACCCTCATCCACATCTACCACATCAGGGGCTACCAACTCTACTTCCGGTTCCACTTCATTTGGTGCTTCCGTAACAGGTTCAACTGCCTCCACTTCATTCGGTGCTTCCGTAACTGGTTCAACCGCTTCCACTTCATTCGGTGCTTCCGTGACTGGTTCAACGGCTTCCACCTTGACTTCCGGCTCCCCATCTGTTTATACCACAACATTAACATATGCAACAACCACAAGCACAGTAGTTGTCTCCTGTTCAGAAACAACTGATTCGAACGGTAACGTCTATACCATTACCACAACCGTACCATGTTCATCTACCACCGCCACTATCACTTCTTGCGATGAGACCGGATGTCATGTAACTACGTCTACCGGTACCGTCGCCACTGAAACCGTTTCTTCCAAATCATACACCACTGTTACCGTCACCCACTGTGACAACAATGGCTGTAACACCAAGACTGTCACTTCTGAATGTCCTGAAGAAACTTCAGCAACTACTACTTCTCCAAAATCATACACTACTGTTACCGTTACTCACTGTGACGACAACGGCTGTAACACTAAGACTGTCACCTCTGAGGCCCCTGAAGCCACAACCACTACTGTTTCTCCAAAGACATACACTACCGCTACTGTTACTCAGTGCGATGACAATGGATGTAGCACCAAGACTGTCACTTCTGAAGCTCCTAAAGAAACTTCAGCAACTACTACTTCTCCAAAATCATACACTACTGTTACCGTTACTCACTGTGACGACAACGGCTGTAACACTAAGACTGTCACCTCTGAGGCCCCTGAAGCCACAACCACTACTGTTTCTCCAAAGACATACACTACCGCTACTGTTACTCAGTGCGATGACAATGGATGTAGCACCAAGACTGTCACTTCTGAAGCTCCTAAAGAAACTTCAGCAACTACTACTTCTCCAAAATCATACACTACTGTTACCGTTACTCACTGTGACGACAACGGCTGTAACACTAAGACTGTCACCTCTGAGGCCCCTGAAGCCACAACCACTACTGTTTCTCCAAAGACATACACTACCGCTACTGTTACTCAGTGCGATGACAATGGATGTAGCACCAAGACTGTCACTTCTGAAGCTCCTAAAGAAACTTCAGCAACTACTACTTCTCCAAAATCATACACTACTGTTACCGTTACTCACTGTGACGACAACGGCTGTAACACTAAGACTGTCACCTCTGAGGCCCCTGAAGCCACAACCACTACTGTTTCTCCAAAGACATACACTACCGCTACTGTTACTCAGTGCGATGACAATGGATGTAGCACCAAGACTGTCACTTCTGAAGCTCCTAAAGAAACTTCAGCAACTACTACTTCTCCAAAATCATACACTACTGTTACCGTTACTCACTGTGACGACAACGGCTGTAACACTAAGACTGTCACCTCTGAGGCCCCTGAAGCCACAACCACTACTGTTTCTCCAAAGACATACACTACCGCTACTGTTACTCAGTGCGATGACAATGGATGTAGCACCAAGACTGTCACTTCTGAAGCTCCTAAAGAAACTTCAGCAACTACTACTTCTCCAAAATCATACACTACTGTTACCGTTACTCACTGTGACGACAACGGCTGTAACACTAAGACTGTCACCTCTGAGGCCCCTGAAGCCACAACCACTACTGTTTCTCCAAAGACATACACTACCGCTACTGTTACTCAGTGCGATGACAATGGATGTAGCACCAAGACTGTCACTTCTGAAGCTCCTAAAGCAACCTCATTGACTACTGCCATTTCCAAGGCTTCTAGTGCAATTTCCACATACTCCAAATCTGCAGCTCCAATAAAGACCTCTACTGGTATCATTGTCCAGTCCGAGGGTATTGCCGCAGGATTGAATGCCAATACTTTGAATGCATTGGTCGGTATTTTCGTTCTTGCTTTCTTTAACTAA

**>*HPF1*_YPS128**

ATGTTCAATCGCTTTAATAAACTTCAAGCCGCTTTGGCTTTGGTCCTTTACTCCCAAAGTGCATTGGGCCAATATTATACCAACAGTTCCTCAATCGCTAGTAACAGCTCCACTGCCGTTTCGTCAACTTCATCAGGTTCTGTTTCCATCAGTAGTTCTATTGTTGAGTCGACCTCATCTGCTTCTGATGTCTCGAGCTCTCTCACTGAGTTAACATCATCCTCCACCGAAGTCTCGAGCACCATTGCTCCATCAACCTCGTCCTCTGAAGTCTCGAGCTCTATTACTTCATCAGGCTCATCAGTCTCCGGCTCATCTTCTATTACTTCATCAGGCTCATCAGTCTCCAGTTCATCTTCTGTCACAGAATCAGGCTCATCCGCCCCAGGTTCATCTACTTCCATTACATCAGGTTCATCTTCTGCCACAGAATCAGGCTCATCAGTCTCCGGTTCATCTACTTCCATTACATCAGGCTCATCCTCCGCCACTGAATCGGGCTCATCAGTCTCCGGTTCAACTTCTGCCACTGAATCAGGCTCATCCGCCTCCGGTTCAACTTCCGCCACTGAATCAGGCTCATCAGTCTCCGGTTCATCTTCTGCCACAGAATCAGGCTCATCAGTCTCCGGTTCATCTACTTCCATTACATCAGGCTCATCCTCCGCCACTGAATCAGGCTCATCCGCCTCCGGTTCAACTTCCGCCACAGAATCAGGCTCATCCGCCTCCGGTTCATCTTCTGCCACTGAATCAGGCTCCGCTTCTTCGGTTCCTAGCTCATCCGGTTCTATCACAGAATCAGGCTCATCCTCATCAGCATCTGAATCATCTATCACACAATCTGGTACCGCTTCCGGTTCATCAGCCTCCAGCACGTCCGGTTCTGTTACACAATCTGGTTCCTCCGTTTCCGGTTCATCAGCTTCTTCTGCTCCAGGTATCTCGAGTTCAATTCCTCAATCAACCTCATCGGCTTCCACTGCCTCCGGTTCTATCACCTCCGGTACCTTAAGTTCTATTACCTCTTCGGCTTCTAGTGCAACTGCAACTGCTTCCAACTCTCTTTCTTCCAGCGATGGTACTATTTATTTGCCTTCTACAACCATCAGTGCTGACATCACACTCACCGGTTCAGTCATTGCAACTGAAGCTGTCGAAGTCGCTGCAGGTGGTAAGTTGACCCTACTTGATGGTGACAAATACGTTTTTTCTGCTGATTTCATAATCCATGGTGGCGTTTTCGTAGAAAAGTCTAAGCCAACTTACCCAGGTACCGAATTCGACATTTCTGGTGAAAACTTTGATGTATCTGGTACCTTTAACGCTGAAGAGCCTGCTGCTTCTTCCGCATCTGCATACTCCTTCACTCCAGGCTCTTTCGATAACAGTGGTGATATTTCTTTGAGTCTATCAGAGTCCACAAAGGGCCAAGTCACATTCTCTCCTTACTCTAACTCTGGTGCTTTCTCTTTCTCAAATGCTATTCTCAATGGTGGTTCCGTCTCTGGTTTGCAACGTAGAGCTGAATCAGGTTCTGTCAACAACGGTGAGATAAATATTGAGAATGGCAGTACCTACGTCGTTGTCGAACCAGTTTCTGGAAGTGGTACAATCAACATCATCTCTGGCAACCTTTACTTGCACTATCCAGACACCTTTACTGGCCAAACTGTTGTATTCAAGGGTGAAGGTGTTCTTGCCGTTGACCCTACCGAAAGCAACACTACCCCTATCCCTGTGGTTGGATACACTGGTGAAAACCAAATCGCCATTACAGCAGATGTAACTGCTCTTTCTTACGACAGTGCTACTGGTGTTTTAACTGCAACACAAGGCAACTCACAATTCTCCTTCTCTATTGGTACTGGATTCTCCAGTTCTGGTTTCAACGTCTCCGAAGGAACATTTGCTGGTGCCTATGCTTATTATCTAAATTACGGAGGTGTTGTTGCTTCCAGCGCTACACCCTCATCCACATCTACCACATCAGGGGCTACCAACTCTACTTCCGGTTCCACTTCATTTGGTGCTTCCGTAACAGGTTCAACTGCCTCCACTTCATTCGGTGCTTCCGTAACTGGTTCAACCGCTTCCACTTCATTCGGTGCTTCCGTGACTGGTTCAACGGCTTCCACCTTGACTTCCGGCTCCCCATCTGTTTATACCACAACATTAACATATGCAACAACCACAAGCACAGTAGTTGTCTCCTGTTCAGAAACAACTGATTCGAACGGTAACGTCTATACCATTACCACAACCGTACCATGTTCATCTACCACCGCCACTATCACTTCTTGCGATGAGACCGGATGTCATGTAACTACGTCTACCGGTACCGTCGCCACTGAAACCGTTTCTTCCAAATCATACACCACTGTTACCGTCACCCACTGTGACAACAATGGCTGTAACACCAAGACTGTCACTTCTGAATGTCCTGAAGAAACTTCAGCAACTACTACTTCTCCAAAATCATACACTACTGTTACCGTTACTCACTGTGACGACAACGGCTGTAACACTAAGACTGTCACCTCTGAGGCCCCTGAAGCCACAACCACTACTGTTTCTCCAAAGACATACACTACCGCTACTGTTACTCAGTGCGATGACAATGGATGTAGCACCAAGACTGTCACTTCTGAAGCTCCTAAAGAAACTTCAGAAACTTCAGAAACCAGTGCTGCCCCTAAGACATACACTACTGCCACTGTTACTCAATGTGATGACAATGGTTGTAACGTCAAGATAATCACCTCTCAAATACCTGAAGCTACTTCAACCGTCACCGCAACTAGTGCTTCTCCAAAGTCATACACTACTGTCACTTCTGAGGGTTCTAAAGCAACCTCATTGACTACTGCCATTTCCAAGGCTTCTAGTGCAATTTCCACATACTCCAAATCTGCAGCTCCAATAAAGACCTCTACTGGTATCATTGTCCAGTCCGAGGGTATTGCCGCAGGTTTGAATGCCAATACTTTGAATGCATTGGTCGGTATTTTCGTTCTTGCTTTCTTTAACTAA
